# Oocyte-stored Mif promotes preimplantation development as an embryonic extracellular signal

**DOI:** 10.64898/2026.11.03.763862

**Authors:** John Xiao He Li, Huasong Wang, Lifang Li, Tursunjan Aziz, Huiying Huang, Chenxin Li, Guangyuan Li, Han Li, Jie Li, Lu Tian, Zihang Wei, Tianyi Zhang, Tong Zhu, Yuling Chen, Haiteng Deng, Xiaotong Wu, Anming Meng

## Abstract

Mammalian preimplantation embryos develop more efficiently in groups than in isolation, suggesting that embryos secrete embryotrophic factors that promote development. However, the endogenous embryotrophic signals responsible for this long-recognized phenomenon remain incompletely defined. Using an unbiased proteomic screen for stable maternal proteins that may function as embryonic extracellular signals, we identified the cytokine macrophage migration inhibitory factor (Mif) as an abundant oocyte-stored protein that persists for several days after fertilization in mice. Quantitative measurements of spent culture medium reveal that developing embryos release Mif, with a substantial fraction derived from the maternal deposit. Extracellular Mif levels peak 3–4 days post fertilization, the same period when *Mif* knockout preimplantation embryos produce its strongest phenotype. Embryos lacking maternal, but not zygotic, *Mif* display defective blastocyst growth even in group culture. Individually cultured embryos exhibit similar blastocyst growth defects that can be rescued by supplementing recombinant Mif at physiologically relevant concentrations. Mif activates the PI3K-Akt-mTORC1 pathway through the cell-surface receptor Cd74 to promote blastocyst growth and cell proliferation. Together, these findings identify Mif as a potential endogenous contributor to the density-dependent embryotrophic effects and reveal that maternally stored proteins may function long after fertilization as extracellular signaling molecules to support preimplantation development in mammals.

## Main text

Although preimplantation mammalian embryos execute an intrinsic developmental program, their growth is critically influenced by extracellular cues from both the maternal reproductive tract and the embryos themselves (1–4). A long-standing observation in reproductive biology is that groups of embryos develop more efficiently than individually cultured embryos in vitro (5–7). Culture in groups or at high embryo density is routinely used in laboratory research (e.g. one embryo per µl of medium, (7, 8)) while culture as individuals or at low embryo density is often favored for human embryos in clinical in vitro fertilization (IVF) (9). This group culture effect suggests that embryos produce diffusible embryotrophic factors that promote development, but the endogenous contributors of these signals remain poorly defined (10–17).

Numerous growth factors, including EGF family ligands, IGF, and FGF, can enhance embryo development when supplemented exogenously (18, 19). However, these candidates were largely selected based on known growth-promoting activity. Mechanistically, the downstream signaling of many growth factors converge to the PI3K-Akt pathway in preimplantation embryos (8, 20). Binding to cell-surface receptors activates PI3K which converts phosphatidylinositol (4,5)-bisphosphate or PI(4,5)P_2_ to (3,4,5)-trisphosphate or PI(3,4,5)P_3_. The phospholipid PI(3,4,5)P_3_ recruits PDK1 and Akt proteins to the plasma membrane by their pleckstrin homology (PH) domains. PDK1 and mTORC2 phosphorylate and activate Akt kinase activity which promotes cell proliferation and suppress apoptosis (21).

Recent proteomic studies on spent culture medium and blastocoel fluid have directly identified numerous proteins released by embryos but distinguishing functional signals from the entire released proteome remains challenging (22–25).

We hypothesized that endogenous embryotrophic factors may include stable proteins inherited from the oocyte, because the oocyte deposits a large reservoir of proteins that supports early development before the robust zygotic control is established (26, 27). Using an unbiased proteomic approach on mouse embryos, we searched for the abundantly expressed maternal proteins that persist after fertilization and could potentially function extracellularly. We favored the embryo over spent medium as the material for proteomics as it offers higher input and allows enrichment of maternal proteins under our experimental design.

This approach identified macrophage migration inhibitory factor (Mif), a pleiotropic cytokine best known for its roles in inflammation but has also been linked to cellular functions including proliferation, apoptosis and motility (28, 29), as a highly abundant oocyte-stored protein that persists into development. Using genetic knockout mice lacking either maternal (mKO) or zygotic *Mif* (zKO) or both (mzKO), we experimentally confirmed that maternal Mif is released by developing embryos and promotes blastocyst growth 3–4 days after fertilization, revealing an unexpected time window of maternal protein function.

## Results

### Identification of Mif in the stable maternal proteome

To identify maternally inherited secreted proteins in mouse embryos, we used α-amanitin to inhibit zygotic genome activation (ZGA) thus preventing interference from the zygotically produced secretome. α-amanitin-treated zygotes could develop up to the two-cell stage when ZGA occurs in mice. We collected treated samples at this stage for proteomics reasoning that the oocyte-stage specific secretome may have further degraded thus lessening its interference with our search for those are stably inherited into the embryonic stages.

Our label-free mass spectrometry-based proteomics data captured 3022 maternal proteins (Fig. 1A, Table S1, and Dataset S1). We screened their UniProt annotations for both conventional (containing signal peptides) and unconventional protein secretion (UPS) (with “extracellular” or “secreted” annotations) (30) and found 210 putative secreted proteins. We further excluded those containing transmembrane domains or lipid anchors or are endoplasmic reticulum-retained and obtained 61 putative soluble proteins. Known secreted proteins Pdcd6ip/Alix and Ppia/Cyclophilin A as well as UPS machinery Hsp90ab1/Hsp90β (31–33) were recovered supporting our screen strategy (Fig. 1A). The cytokine Mif, a leaderless protein that is secreted by regulated UPS (34, 35), ranked the 7^th^ among the 61 proteins and the 126^th^ among all proteins by abundance (Fig. 1A). Mif was also highly abundant among the maternal proteome in zygotes according to our re-analysis of published proteomic data (27), ranking even higher (292^nd^/4045) than the abundant subcortical maternal complex proteins (SCMC), Zbed3 and Nlrp2 (26) (Fig. 1B). SCMC components as well as Mif remained abundant from zygote to the two-cell stage while Sfn/14-3-3σ, Oosp1 and Gdf9, which are mainly secreted by the oocyte (36–38), dropped in ranking (Fig. 1, A and B). Our sample collection strategy could enrich stable maternal proteins.

**Figure 1.**
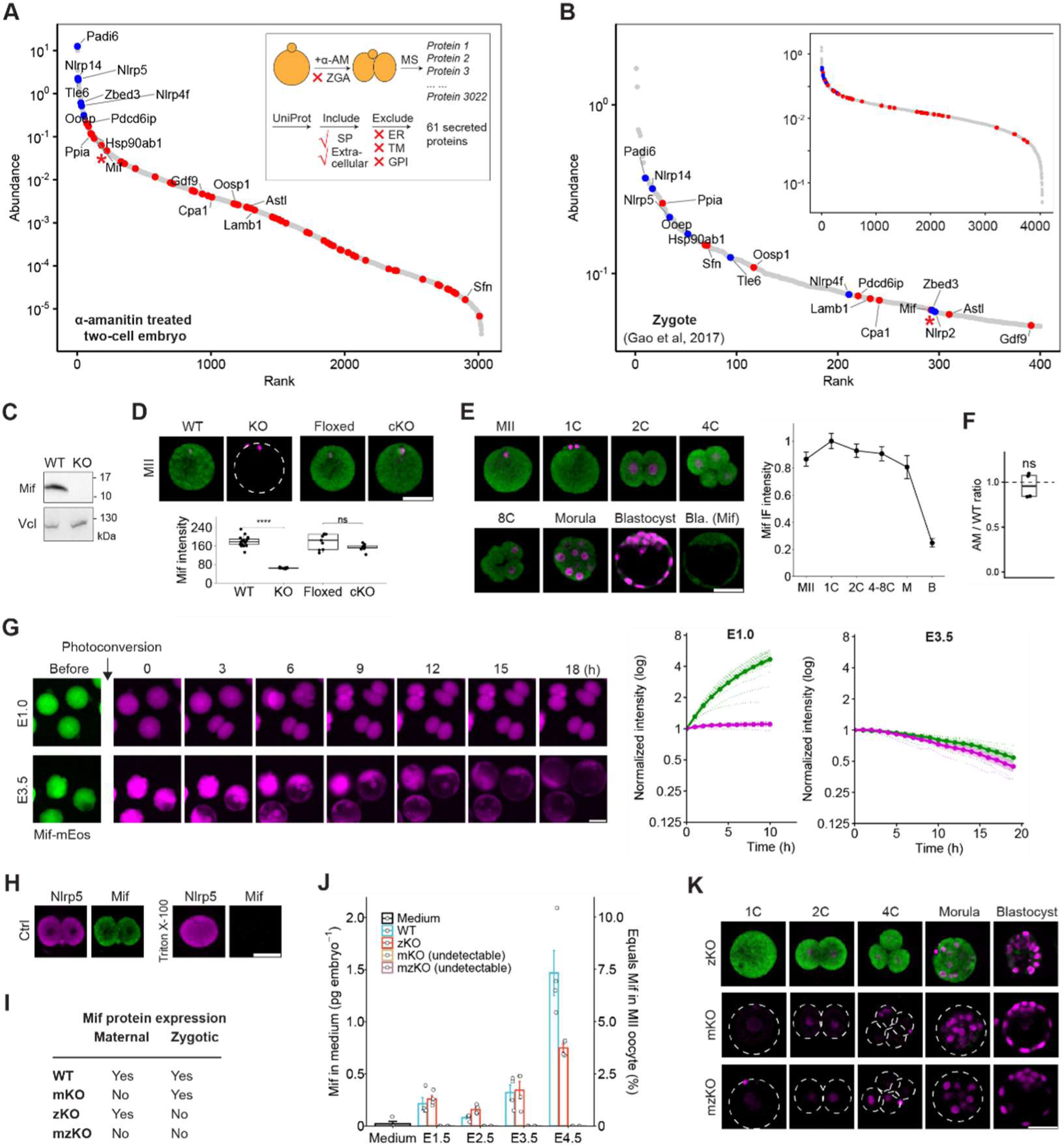
The developing embryo releases maternal Mif protein. **A:** Relative protein abundance in mouse two-cell embryos treated with α-amanitin (α-AM) from zygotes to suppress zygotic genome activation (ZGA) which happens at the two-cell stage in mice. Inset shows sample collection and screen strategy. Mass spectrometry (MS)-identified proteins were screened for signal peptide (SP) or extracellular/secreted annotations in UniProt database. Proteins with endoplasmic reticulum (ER) localization, transmembrane domains (TM) or lipid anchors (GPI) were excluded. Putative secreted proteins (red), subcortical maternal complex (blue), and Mif (asterisk) are labeled. **B:** Relative protein abundance of the top 400 proteins in mouse zygotes re-analyzed from published data (27). Inset shows all proteins. Overlapped proteins with (A) are labeled in color. **C:** Validation of Mif antibody by Western blot of *Mif*^+/+^ (WT) and *Mif*^-/-^ (KO) liver extracts. Vinculin (Vcl), loading control. **D:** (D and E) Immunofluorescence of Mif (green) and 4’,6-diamidino-2-phenylindole (DAPI) (magenta). MII oocytes from WT, KO, *Mif^fl/fl^* (Floxed), and *Zp3-Cre;Mif^fl/fl^* (cKO) mice. The boxplot shows median and interquartile range (IQR) with whiskers marking the inlier range; N = 20, 12, 6, and 6; two-sided Welch’s t-test with Holm correction. **E:** Developmental stages. Zygote (1C) is at Embryonic day (E) 0.5; two-cell stage (2C), E1.5; four-cell and eight-cell stages (4C/8C), E2.5; morula (M), E3.5; blastocyst (B), E4.5. The plot shows mean ± s.e.m. level normalized to zygote stage of WT (as 1) and KO (as 0); N = 9, 4, 5, 3, 5, and 2. **F:** Zygotes were treated with α-amanitin as in (A). The plot shows treated versus non-treated (WT) ratio of Mif immunofluorescence intensity in each experiment; N = 4; two-sided Student’s paired t-test. **G:** Mif protein turnover measured by photoconversion pulse-chase assay. An mRNA encoding for Mif-mEos fusion protein was injected at E0.5. A fraction of Mif-mEos (green) was photoconverted (magenta) at E1.0 and 3.5, respectively, and imaged over time. Note that the continuous translation of mRNA led to increase of unconverted Mif-mEos at E1.0; the image intensity was adjusted for contrast independently between E1.0 and 3.5 and thus incomparable. The plots show mean (solid line) ± s.e.m. (shade) of normalized fluorescence intensity starting 2 h post photoconversion of individual embryos (dotted lines); N = 19 and 5. **H:** Immunofluorescence of insoluble protein Nlrp5 (magenta) and Mif (green). Note that detergent extraction by 0.1% Triton X-100 released all intracellular Mif. **I:** Summary of Mif protein expression in KO. **J:** Quantification of secreted Mif protein in the spent culture medium by ELISA (left axis) and its equivalence to MII oocyte Mif level (right axis). Fresh medium (N = 4) was a negative control. Culture started at E0.5 and spent medium was collected daily at the indicated time points after removing embryos. The plot shows mean ± s.e.m. per embryo; WT (N = 4); zygotic KO (zKO, from *Zp3-Cre;Mif^fl/fl^* ♀ × *Mif^-l-^* ♂ parents, see Table S3, N = 4); maternal KO (mKO, *Mif^-l-^* ♀ × WT ♂, N = 3); maternal-zygotic KO (mzKO, *Mif^-l-^* ♀ × *Mif^-l-^* ♂, N = 2). Note that mKO and mzKO levels were undetectable and shown as zero. **K:** Immunofluorescence of Mif (green) and DAPI (magenta). Scale bar, 50 μm.

We verified the expression of maternal Mif protein using a specific antibody that we validated by *Mif* genetic knockout (KO) mice (Fig. 1, C and D, and Fig. S1A). Immunofluorescence showed that Mif level remained largely unchanged from MII oocyte through morula stages but dropped by 75% in blastocysts (Fig. 1E). In contrast, the Mif mRNA level was low in MII oocytes and zygotes and gradually increased after ZGA; mRNA translation activity was tightly coupled to its transcription at the embryonic stages, according to the published RNA-seq and ribo-seq data (39) (Fig. S1B). Further, Mif protein remained at the same level in the absence of ZGA in α-amanitin-treated two-cell embryos (Fig. 1F). Thus, the abundant Mif protein at the early embryonic stages may be inherited as stable maternal proteins.

To investigate the stability of maternal Mif protein, we modeled the turnover of endogenous Mif with the exogenously expressed Mif that is fused to a widely used photoconvertible fluorescent protein mEos (40). Upon short ultraviolet illumination (pulse labeling), a fraction of Mif-mEos would turn red from green creating a finite pool for a chase assay. By time-lapse imaging of converted proteins, we observed drastically different Mif behaviors at E1.0 and E3.5. The converted protein level remained change at E1.0, while it was removed at E3.5 with an apparent half-life of 17.4 h (Fig. 1G). Overall, Mif protein was sustained at high levels in developing embryos several days after fertilization.

### Timed release of maternal Mif by the embryo

Next, we used ELISA to detect if Mif is released by the developing embryos. We collected the spent culture medium every day between embryonic day (E) 0.5 and 4.5 which is the entire span of the preimplantation development. Embryos were cultured in groups as Mif levels from individual culture were below the ELISA detection limit. We failed to detect Mif protein in the fresh medium (Fig. 1J and Table S2). Further, the Mif level released by the mzKO embryos was undetectable suggesting that this assay was Mif-specific (Fig. 1J). The secretion activity of WT embryos started low right after fertilization but increased by about 6-fold between day 3–4 during the morula and blastocyst stages. We could not measure Mif released during fertilization but it is supposed to be at a low level as the intracellular Mif protein levels remains nearly unchanged from MII oocytes to zygotes (Fig. 1E). Total released Mif per embryo between E0.5–4.5 was 2.1 ± 0.2 pg (mean ± s.e.m., N = 4) (Fig. 1J, left axis). Using a detergent extraction condition that released all intracellular Mif (Fig. 1H), we determined that each MII oocyte contained 20.4 ± 1.2 pg maternal Mif protein by ELISA, with results from two separate experiments well matched (mean ± s.e.m., N = 4, Table S2) and agreeing relatively well with that from quantitative Western blot (Fig. S1G). Thus, total released Mif equals about 10% of Mif in the oocyte (Fig. 1J, right axis). A significant amount of Mif is released by the embryo.

To determine the relative contribution of the maternal and zygotic sources for released Mif, we compared the Mif levels released by WT with zKO that lacks both zygotic *Mif* alleles. Traditionally, zKO was generated by crossing *Mif*^+/-^ parents. However, the mixed genotypes in each litter are incompatible with group culture for measuring released Mif. Instead, we used the *Zp3-Cre;Mif^fl/fl^* female mice carrying an oocyte-specific *Cre* and two floxed *Mif* alleles (Fig. S1A). Despite the efficient Cre-mediated excision of the entire *Mif* gene, we failed to detect obvious Mif protein level changes in their oocytes (cKO) compared to *Mif^fl/fl^* (Floxed) oocytes indicating an intact maternal Mif protein deposit (Fig. 1D). Crossing *Zp3-Cre;Mif^fl/fl^* females with *Mif^-l-^* males gave embryos that were all zKO (Table S3). Maternal Mif protein persisted through the morula stage in zKO (Fig. 1, I and K). The Mif released by zKO in group culture is supposed to be mainly from the maternal source, which sums to 1.5 ± 0.1 pg between E0.5–4.5 (mean ± s.e.m., N = 4) (Fig. 1J). The difference in released Mif levels between WT and zKO embryo was mainly at the E3.5–4.5, which is most likely due to the lack of zygotic contribution and coincides with the zygotic Mif expression window (Fig. S1B). By subtracting 1.5 pg (zKO) from 2.1 pg (WT) total released Mif, we deduced that 1.5 pg (70%) and 0.6 pg (30%) were possibly released from the maternal and zygotic sources, respectively (Fig. S1H).

We verified the maternal contribution by measuring Mif released by mKO embryos. Because Mif protein remained in the cKO oocyte, we used germline KO females to completely deplete maternal Mif and crossed *Mif^-l-^* ♀ with WT ♂ to generate mKO (Table S3). The Mif level released from these mKO embryos was undetectable (Fig. 1J). These results together suggest that developing embryos release Mif, possibly with the maternal deposit providing the substantial fraction.

### Maternal Mif promotes blastocyst development

We tested if Mif could contribute to the density-dependent embryotrophic effects. We set up high density (HD) and low density (LD) culture which were 20 embryos in 20 μl medium or 1 μl^-1^ and one embryo in 30 μl or 0.03 μl^-1^, respectively. The 20 μl volume at HD is picked to match the average embryo number of superovulated female mice. The 30 μl volume at LD safely exceeds the 25 μl volume limit to concentrate embryotrophic factors and is expected to inhibit development (5, 6). We switched from HD to LD between E3.5 and E4.5 stages instead of LD throughout to avoid accumulative effects due to dilution of known embryotrophic factors at earlier stages (17) (Fig. 2A). This period also coincides with the main period of Mif release (Fig. 1J). During this period, released Mif concentration in the spent culture medium at LD was 21.9 ± 5.3 pg ml^-1^ (mean ± s.e.m., N = 3), 50 times lower than that at HD which was 1090 ± 139 pg ml^-1^ (N = 2) (Fig. 2B). The total Mif released by each embryo was, however, similar between LD and HD (Fig. 2B). Low density culture dilutes the concentration of released Mif.

**Figure 2.**
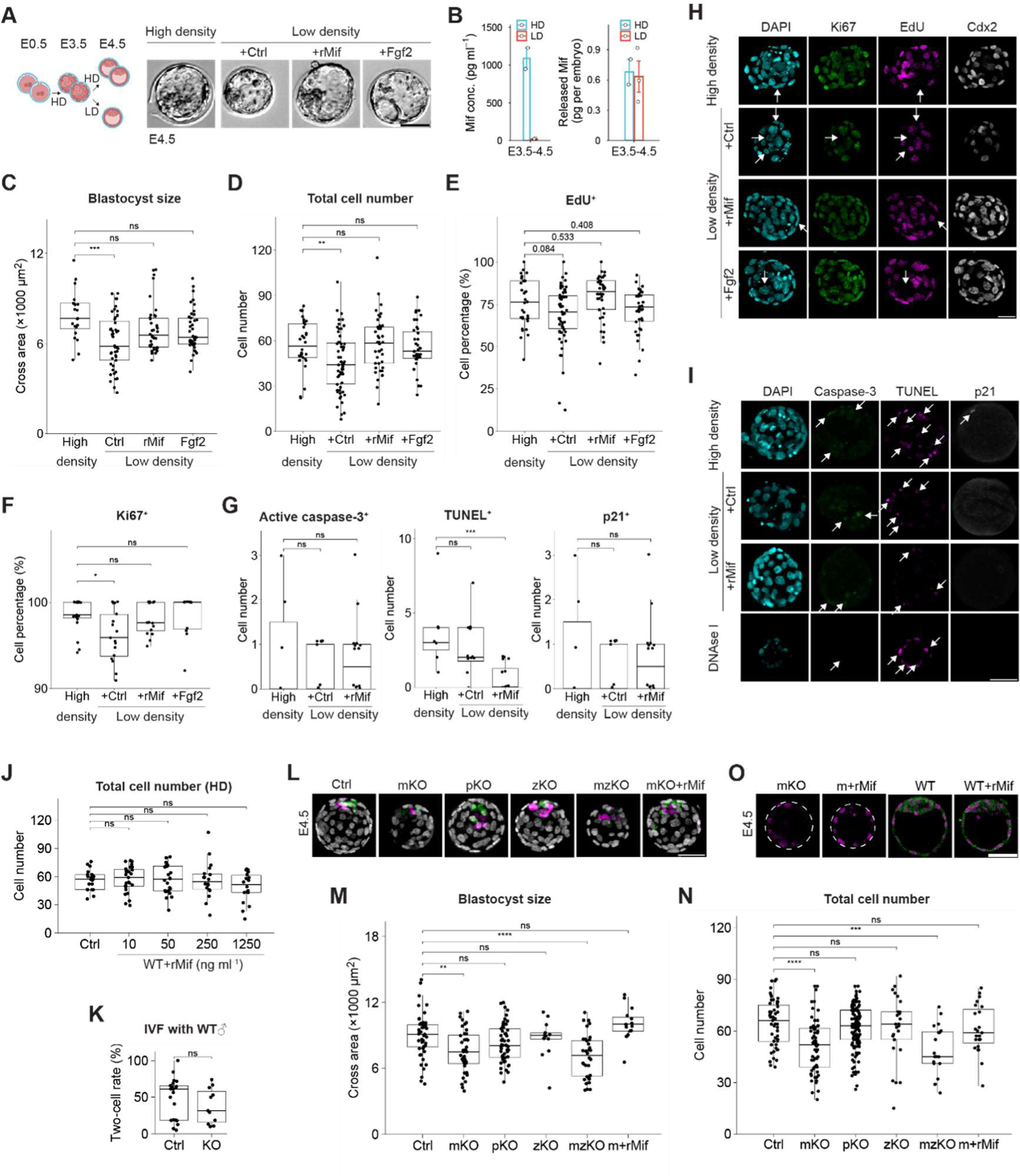
Extracellular Mif promotes blastocyst development. **A:** (A–I) WT zygotes were cultured at high density (HD; 20 embryos in 20 μl; 1 μl^-1^) E0.5–3.5; compact morulae were washed and cultured at HD or low density (LD; 1 in 30 μl; 0.03 μl^-1^) between E3.5–4.5; 100 pg ml^-1^ boiled (Ctrl) or normal recombinant Mif (rMif) or 40 μg ml^-1^ Fgf2 (41) was added to rescue LD effect; E4.5 blastocysts were examined. (A) The images show live blastocysts. **B:** Quantification of Mif concentration in the spent culture medium and total released Mif per embryo E3.5–4.5 by ELISA. The plot shows mean ± s.e.m.; N = 2 and 3; LD was measured by combining media of 10 individually cultured embryos (see Methods). **C:** Cross-sectional area. N = 20, 47, 40, and 40; Dunnett’s test. **D:** (D and E) Total cell number and EdU^+^ incorporation (cell proliferation marker). N = 32, 59, 42, and 39; Dunnett’s test. **F:** Proliferating Ki67^+^ cell percentage. N = 19, 16, 16, and 11; Dunnett’s test. **G:** Apoptotic (TUNEL^+^ and active-caspase-3^+^) and p21^+^ cell number. N = 7, 12, and 16; Dunnett’s test. **H:** Immunofluorescence of DAPI (cyan), Ki67 (green), EdU (magenta), and Cdx2 (gray, trophectoderm marker) (maximum z-projection). Arrow, cells negative for each marker. **I:** Immunofluorescence of DAPI (cyan), active caspase-3 (green), TUNEL (magenta), and p21 (gray) (maximum z-projection). Fixed blastocysts were treated with DNAse I as a positive control for TUNEL (single z-slice image). Arrow, cells positive for each marker. **J:** WT zygotes at HD were treated with 250 ng ml^-1^ boiled rMif (Ctrl) or indicated concentration of normal rMif E0.5–4.5. The plot shows total cell number; N = 21, 27, 24, 18, and 18; Dunnett’s test. **K:** Oocyte competence assay. *Mif^+l+^* and *Mif^-l-^* cumulus-oocyte complexes (COCs) underwent in vitro fertilization (IVF) with WT sperm. N = 22 and 11 females with a total of 534 and 255 COCs, respectively; two-sided Welch’s t-test. **L:** (L–N) *Mif* KO zygotes were cultured at HD E0.5–4.5; E4.5 blastocysts were examined. Immunofluorescence of Cdx2 (white), Sox2 (green), and Sox17 (magenta) (maximum z-projection). Ctrl (*Mif^+l-^* ♀ × WT ♂); pKO (paternal KO, WT ♀ × *Mif^-l-^* ♂, reciprocal cross of mKO); m+rMif (mKO treated with 6.3 ng ml^-1^ rMif; see Methods). **M:** Cross-sectional area. N = 47, 39, 52, 13, 34, and 15; Dunnett’s test. **N:** Total cell number. N = 59, 63, 137, 25, 19, and 23; Dunnett’s test. **O:** Zygotes were treated with 200 ng ml^-1^ rMif E0.5–4.5. Immunofluorescence of Mif (green) and DAPI (magenta). Note that intracellular Mif was absent in mKO and did not increase due to rMif treatment. Scale bar, 50 μm.

Although starting from the same batch of morulae, LD culture resulted in smaller blastocysts with fewer cells at E4.5 (Fig. 2, C and D, HD versus LD +Ctrl). The rate of blastocyst formation was slightly decreased suggesting that LD mainly affected blastocyst expansion instead of the initiation of cavitation (Fig. S2A). Supplementing a known embryotrophic factor Fgf2 (41) rescued these phenotypes at LD culture (Fig. 2, C and D, LD +Ctrl versus +Fgf2). Supplementing as low as 100 pg ml^-1^ rMif, but not its boiled and inactive form, also rescued blastocyst size and cell number at LD culture (Fig. 2, C and D, LD +Ctrl versus +rMif). rMif rescued both trophectoderm and inner cell mass cell numbers (Fig. S2B). We used 100 pg ml^-1^ to achieve a moderate, ∼6-fold increase from physiological extracellular Mif concentration at LD of 21.9 pg ml^-1^ (Fig. 2B). In contrast, rMif at concentrations that span several magnitudes higher than that of accumulative extracellular Mif concentration (see Methods) had minimal effects on embryos at HD, even if treated throughout E0.5–4.5 (Fig. 2J and Fig. S2C). Because the effective concentration of Mif is very low, the embryo-derived Mif concentration at HD culture has already surpassed the effective concentration. In such, exogenous supplementation has no further effects on embryos at HD. Overall, exogenously supplemented extracellular Mif could promote blastocyst development at LD culture.

To determine whether LD reduced cell number by increasing apoptosis (6, 8, 20) or decreasing cell proliferation, we examined the markers for both cell proliferation (Ki67 staining (42) and pulse 5-ethynyl-2’-deoxyuridine/EdU incorporation (43)) and apoptosis (active caspase-3 staining (44) and TUNEL assay (10)). Both the Ki67^+^ and EdU^+^ cell percentages were downregulated at LD but could be rescued by rMif supplementation (Fig. 2, E, F and H, and Fig. S2B). Surprisingly, switching from HD to LD at E3.5 did not markedly increase active caspase-3 staining and TUNEL staining in blastocysts (Fig. 2, G and I). Cdkn1a/p21, a marker for p53 activation (45), was also unchanged at LD (Fig. 2, G and I). The effect of Mif during the blastocyst stage is mostly promoting cell proliferation.

We then checked the function of endogenous Mif by examining the development of mKO, zKO, and mzKO embryos at HD culture. We also examined the reciprocal cross of mKO (pKO, WT ♀ × *Mif^-l-^* ♂) to control for the missing zygotic allele (Table S3). As the KO oocytes had unchanged developmental rate to the two-cell stage (46), any mKO phenotype is most likely due to the disruption of post-fertilization functions of maternal Mif instead of long-lasting effects of oocyte immaturity (Fig. 2K). Beyond the two-cell stage, mKO and mzKO, but not pKO and zKO, impaired embryo development. Their strongest phenotype was at E3.5 and 4.5 stages as a lower blastocyst rate (Fig. S2D). The cavitated mKO and mzKO, but not pKO and zKO, blastocysts had a smaller size (Fig. 2, L and M). We also found a reduced cell number in mKO and mzKO blastocysts, which was more sensitive to the embryo density than blastocyst size (Fig. 2N and Fig. S2E). Furthermore, EdU^+^ cell percentage was not statistically different in mzKO suggesting either very subtle changes in cell cycle control or additional mechanisms which lead to arrested growth (Fig. S2, F and G). These phenotypes caused by the lack of the maternal but not zygotic alleles coincided with the dominant contribution from the maternal pool for released Mif (Fig. 1J). Importantly, all phenotypes of mKO could be rescued by rMif at a physiologically relevant concentration (see Methods) indicating that the defects were specifically due to loss of Mif (Fig. 2, L–N, and Fig. S2, D and E). Depleting the endogenous maternal Mif could block blastocyst development even at HD culture.

It has also been reported that Mif participates in the intracellular signaling to suppress fibroblast cell proliferation (47). However, rMif treatment did not markedly increase the intracellular Mif level in either WT or mKO embryos thus favoring an extracellular mode of action (Fig. 2O). In sum, rMif in the medium can substitute the endogenous Mif to drive blastocyst expansion by increasing cell proliferation. Thus, the endogenous Mif most likely functions as an extracellular, embryotrophic factor.

### Embryonic expression and function of Mif receptors

Besides its cognate receptor Cd74, Mif may also activates co-receptors Cd44, Cxcr2, Cxcr4, and Ackr3/Cxcr7 (28, 29). Transcriptomic data shows that embryos express *Cd74*, *Cxcr4*, and *Ackr3* (Fig. S3A). By Western blot and immunofluorescence, we verified Cd74 expression was strictly zygotic (Fig. 3A) while Cxcr4 was also maternally expressed in MII oocytes (Fig. S4A). Cd74 protein was first detected at the eight-cell stage with its level peaking in morulae and blastocysts (Fig. 3B and Fig. S3B). The majority of Cd74 was cytoplasmic and nuclear (48), which is consistent with its distribution in the somatic cell types: only 2–5% of Cd74 is on the cell surface (49). Cd74 was also enriched at cell-cell junctions in morulae when overexpressed (Fig. 3C and Fig. S3B). Endogenous Cxcr4 was localized throughout cytoplasm while exogenously expressed Cxcr4 was enriched on the plasma membrane (Fig. S4, A and B). We failed to detect Cd44 or Ackr3 by immunofluorescence in mouse early embryos at any stage. At least, cell-surface Mif receptors Cd74 and Cxcr4 are expressed in embryos.

**Figure 3.**
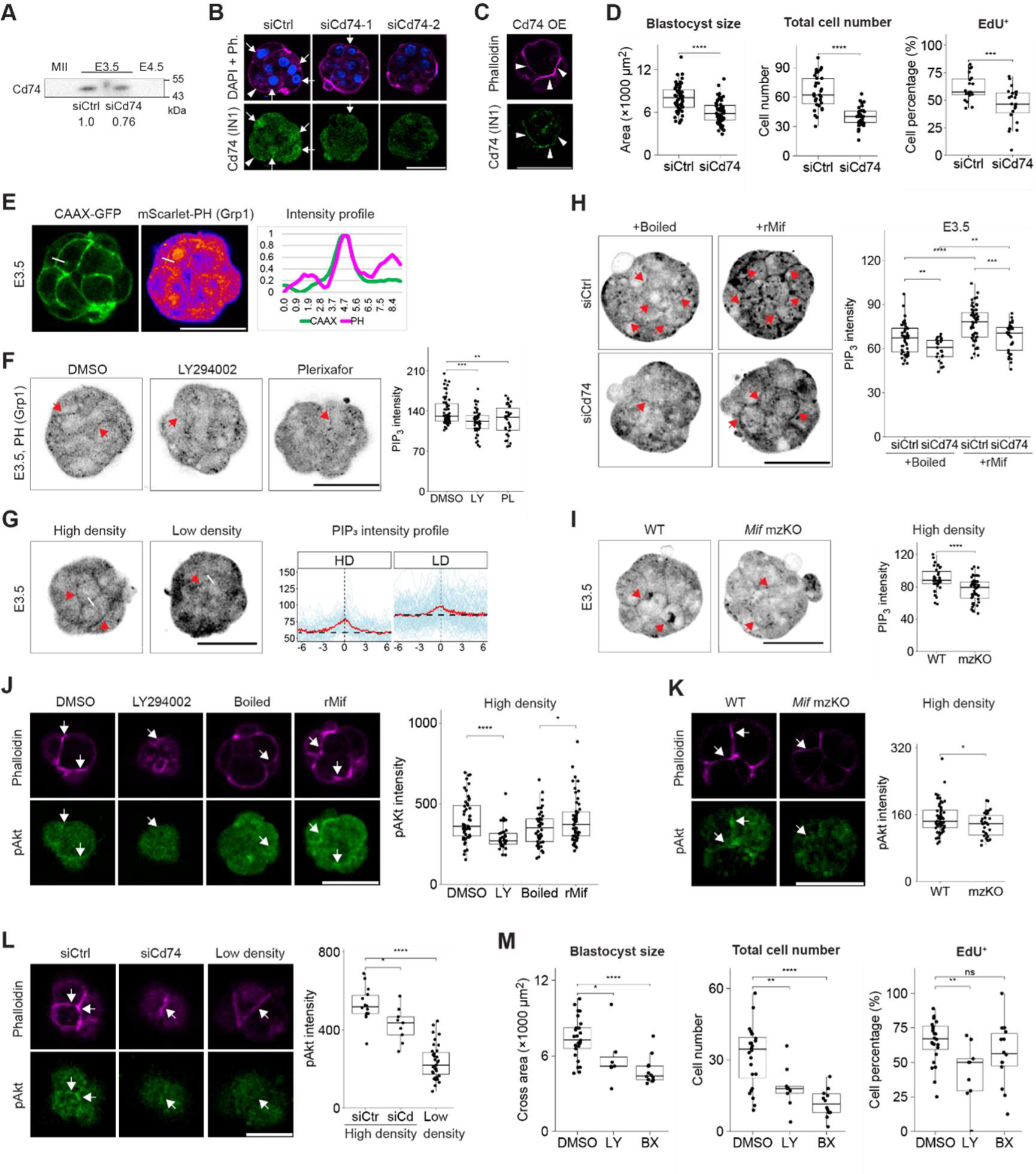
Mif activates PI3K-Akt signaling through the cell-surface Cd74 receptor. **A:** Western blot of 45 oocytes or embryos by anti-Cd74 (clone LN-2). **B:** (B and C) Immunofluorescence of Cd74 (clone IN1) (green), phalloidin (magenta), and DAPI (blue) in E3.5 morulae. siRNA-1 and −2 cause weaker and stronger depletion, respectively. Arrow and arrowhead, Cd74 in the plasma membrane and nuclear regions, respectively. **C:** Cd74 mRNA was injected into zygotes for overexpression (OE). Arrowhead, junctional localization. **D:** siCd74 E4.5 blastocysts at HD culture. N = 58 and 59 (for size); 31 and 31 (for cell number and percentage); two-sided Welch’s t-test. **E:** Live image of plasma membrane marker CAAX-GFP and PI(3,4,5)P_3_ sensor mScarlet-PH (GRP1) in morulae; sensors were expressed from mRNAs injected in zygotes. Intensity across cell-cell junction along the white line was plotted on the right. **F:** (F–I) Live images of PI(3,4,5)P_3_ sensor (inverted grayscale) at junctions (arrow) in morulae; the boxplots show junction intensity. Morulae were treated with 10 μM LY294002 (PI3K inhibitor) and 100 nM Plerixafor (Cxcr4 inhibitor) for 1 h. N = 58, 45, and 34; Dunnett’s test. **G:** Morulae were cultured at HD or LD E3.0–3.5. Intensity across the junction (white lines in images for example) was measured. The plot shows individual intensity profile (light blue) and their average (red) and baseline (dash); N = 49 and 86; note that y-axis starts from 50 artificial units. Total PI(3,4,5)P_3_ intensity at LD culture was higher possibly due to slower sensor degradation in the arrested embryos but its junctional enrichment (elevation between red and dash lines) was weaker. **H:** Morulae treated with 100 pg ml^-1^ rMif for 5 min. N = 51, 25, 53, and 33; two-sided Welch’s t-test with Holm correction. **I:** *Mif* mzKO at HD culture. N = 35 and 48; two-sided Welch’s t-test. **J:** (J–L) Immunofluorescence of phospho-Akt S473 (green) and phalloidin (magenta) at apical junctions (arrow) in E3.5 morulae; the boxplots show junction intensity. Morulae were treated with 25 μM LY294002 for 30 min or 100 pg ml^-1^ rMif for 20 min. N = 50, 31, 44, and 51; two-sided Welch’s t-test with Holm correction. **K:** *Mif* mzKO at HD culture. N = 57 and 31; two-sided Welch’s t-test. **L:** siCtrl and siCd74 were cultured at HD and WT were at LD E3.0–3.5. N = 14, 9, and 35; Dunnett’s test. **M:** WT morulae at HD culture were treated with DMSO, 10 μM LY294002 or 5 μM BX795 (BX, PDK1 inhibitor) E3.5–4.0. N = 25, 7, and 13 (for size); 9, 13, and 12 (for cell number and percentage); Dunnett’s test. Scale bar, 50 μm (except for K, 25 μm).

We examined the zygotic Cd74 function by knockdown (KD) with small interfering RNAs (siRNAs). Immunofluorescence confirmed KD by two previously validated siRNAs injected into zygotes (Fig. 3B). siRNA-1 (50) partially blocked blastocyst formation (Fig. S3F) while siRNA-2 (51) arrested all embryos at the morula stage. Because *Cd74* genetic KO did not completely block live birth in mice (52), We used the more phenotypically related siRNA-1 for all experiments despite it only partially depleted Cd74 proteins by 24% (Fig. 3A). Nonetheless, Cd74 KD decreased blastocyst size and cell number at HD culture, more strongly than *Mif* KO (Fig. 3D). Temporal analysis of cell numbers showed that the reduction first emerged at E3.5 in Cd74 KD, without recovering even if continued to be cultured to E5.5 (Fig. S3C). Outgrowth assay revealed that Cd74 KD disrupted postimplantation development in vitro (Fig. S3H). Treating embryos with inhibitors for Cxcr4, but not Cxcr2 or Ackr3, slightly decreased blastocyst cell number (Fig. S4C). Cd74 may be required for blastocyst development with Cxcr4 playing a contributing role.

Mechanistically, Cd74 KD suppressed cell proliferation without overtly increasing apoptotic cells (Fig. 3D and Fig. S3, E and G). Further, Cd74 KD partially blocked the rescue effects of blastocyst cell numbers by rMif at LD culture (Fig. S3D). Overall, extracellular Mif likely depends on the cell-surface receptor Cd74 to mediate its embryotrophic functions.

### Mif induces PI(3,4,5)P_3_ signaling activity through the Cd74 receptor

The PI3K-Akt signaling pathway is widely recognized and extensively studied as a downstream response to the embryotrophic factors (8, 20). We tested if extracellular Mif could activate the PI3K-Akt pathway at the morula stage when the KO phenotypes start to emerge. Using a fluorescent PH domain sensor, we showed that PI(3,4,5)P_3_ was enriched at the cell-cell junctions in live embryos (8) (Fig. 3E). This junctional PI(3,4,5)P_3_ signal was specific because of its sensitivity to the PI3K inhibitor, LY294002 (Fig. 3F). It was relatively depleted from the junction at LD culture (Fig. 3G). rMif treatment at the physiological concentration strongly upregulated junctional PI(3,4,5)P_3_ (Fig. 3H, siCtrl +Boiled versus +rMif). Importantly, this activating effect of rMif was partially blocked by Cd74 KD and an anti-Cd74 neutralizing antibody (clone LN-2) (53), consistent with Cd74’s junctional localization (Fig. 3H and Fig. S5A). Similarly, junctional PI(3,4,5)P_3_ was depleted by *Mif* mzKO and Cxcr4 inhibitor Plerixafor (Fig. 3, F and I). Using a specific antibody of Akt S473 phosphorylation as a direct readout of Akt activation, we showed that pAkt was also enriched at junctions in morulae. Its localization was sensitive to LY294002 thus placing Akt downstream of PI3K (Fig. 3J). Like PI(3,4,5)P_3_, junctional pAkt enrichment was promoted by rMif treatment but inhibited by *Mif* mzKO, Cd74 KD, and LD culture (Fig. 3, J–L). In contrast, the majority of Akt protein was cytoplasmic (Fig. S5B). Overall, extracellular Mif activates the PI3K-Akt pathway possibly through Cd74 as its cell-surface receptor, with Cxcr4 as one of the potential co-receptors.

Furthermore, we tested the function of PI3K-Akt pathway in development using pharmacological inhibitors. Treating morulae at HD with low doses of LY294002, PDK1 inhibitor BX795 (54), and Akt inhibitor MK2206 (55) severely inhibited blastocyst growth with MK2206 causing a stronger phenotype of complete morula death (Fig. 3M and Fig. S6A). These inhibitors suppressed cell proliferation while activated apoptosis (Fig. S6, B and F). In addition, BX795 completely blocked the rescue effect of rMif on blastocyst cell number at LD culture (Fig. S6C). Extracellular Mif likely requires PI3K-PDK1-Akt pathway activity to promote blastocyst development.

### Mif–Cd74 axis maintains mTORC1 pathway activity

To understand the underlying mechanisms of extracellular Mif function, we therefore performed bulk RNA-seq on E4.0 blastocysts of 6 conditions as follows (Fig. 4A). Four conditions were cultured at HD (*Mif* WT, *Mif* mzKO, siCtrl, and siCd74). In the other two conditions (Boiled and rMif), WT embryos were switched from HD to LD at E3.5 with the supplementation of 100 pg ml^-1^ boiled or normal rMif and collected 12 h later. “Boiled” also serves as the baseline condition of LD for comparison with “siCtrl” as the baseline condition of HD.

**Figure 4.**
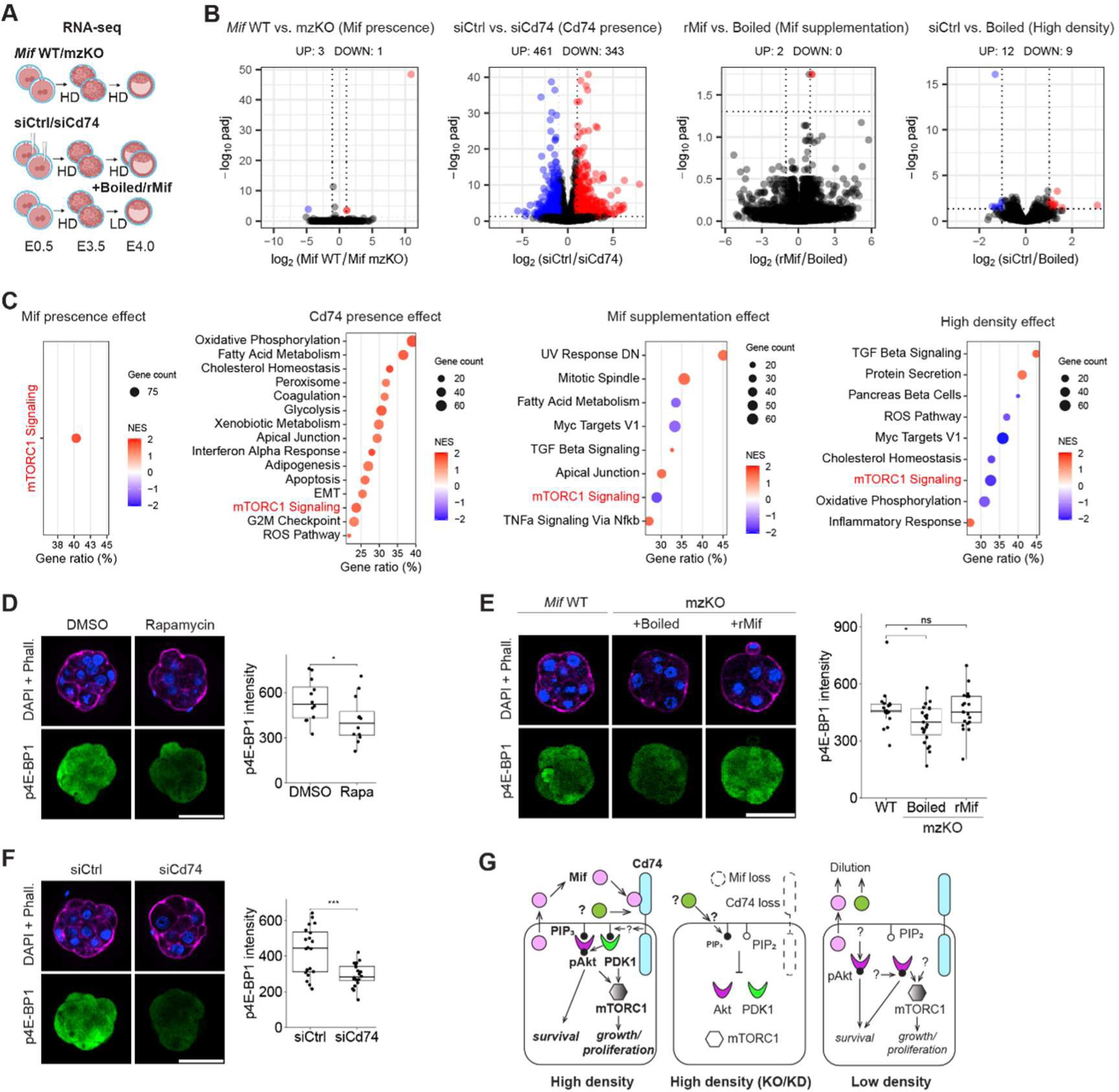
Mif–Cd74 axis maintains mTORC1 activity. **A:** Experimental design of RNA-seq (N = 5 replicates for *Mif* WT and 4 for the rest). siRNAs were injected at E0.5; embryos were cultured at HD or LD from E3.5 with the supplementation of 100 pg ml^-1^ rMif; blastocysts were collected for bulk poly(A) RNA-sequencing at E4.0. **B:** Volcano plot with counts of differentially expressed genes (DEG). x-axis, log_2_ fold change (LFC); y-axis, -log_10_ adjusted p-value. Dash lines, |LFC| ≥ 1 and adjusted p < 0.05. **C:** Hallmark pathways identified by Gene Set Enrichment Analysis (GSEA) (adjusted p < 0.05). Gene count is of the core enrichment genes; gene ratio = core enrichment gene count/gene set size; NES, normalized enrichment score. The common pathway in all four comparisons is in red (mTORC1 Signaling). ROS, reactive oxygen species; EMT, epithelial-mesenchymal transition. **D:** (D–F) Immunofluorescence of DAPI (blue), phalloidin (magenta), and phospho-4E-BP1 T37/46 (green). E3.5 morulae were cultured at HD with 100 nM rapamycin for 30 min. N = 12 and 12; two-sided Welch’s t-test. **E:** E3.5 mzKO morulae were cultured at HD with 6.3 ng ml^-1^ rMif for 6 h (same concentration as in Fig. 2, L–N). N = 18, 23, and 20; Dunnett’s test. **F:** E3.5 morulae at HD culture. N = 20 and 20; two-sided Welch’s t-test. **G:** Model. Embryo-derived Mif (pink circle) binds to Cd74 (cyan) to maintain PI3K-Akt-mTORC1 pathway activity and promote cell growth and proliferation, which is disrupted by genetic depletion of *Mif* (KO) or *Cd74* (KD) even when cultured at HD. At LD culture, Mif is diluted and PIP_3_ (black dot) generation is inhibited; yet, Akt and mTORC1 activation is maintained by an unknown mechanism which prevents complete developmental arrest and may involve other potential extracellular factors (olive circle) (see Discussion).

We focused on four comparisons: *Mif* WT versus mzKO (Mif presence effect), siCtrl versus siCd74 (Cd74 presence effect), rMif versus Boiled (Mif supplementation effect), and siCtrl versus Boiled (High density effect). Note that the directions of these comparisons showed concordant phenotypic changes; for example, the first conditions all promote cell proliferation compared to the second conditions. We asked if these effects also brought similar transcriptional changes. Except for “Cd74 presence”, there were a few differentially expressed genes (DEG) (Fig. 4B, Fig. S7B, and Dataset S2). Although HD culture and Mif treatment could induce strong signaling responses (Fig. 3), it seems that they induced only subtle transcriptomic changes. Nevertheless, we identified mTORC1 (mechanistic target of rapamycin complex 1) signaling pathway as enriched in all four comparisons using Gene Set Enrichment Analysis (GSEA) on the well-established Hallmark Pathway gene sets (56, 57) (Fig. 4C and Dataset S3). Meanwhile, 7 other pathways enriched in at least two comparisons were Myc targets (version 1), TGF-β signaling, apical junction, cholesterol homeostasis, fatty acid metabolism, oxidative phosphorylation, and reactive oxygen species.

mTORC1 pathway is a major downstream route of PI3K-Akt signaling. Akt indirectly activates mTORC1 to phosphorylate S6K1 and 4E-BP1, which promotes cap-dependent translation of proteins. mTORC1 integrates growth factor signals to promote biomass synthesis for cell growth and proliferation (58). The positive normalized enrichment score (NES) of the mTORC1 signaling gene set in “Mif presence” and “Cd74 presence” comparisons suggest pathway downregulation in the depletion conditions (Fig. 4C). We validated the mTORC1 activity with phospho-4E-BP1 staining as a direct readout. In morulae, p4E-BP1 level was sensitive to the inhibition of mTORC1 by acute, low-dose rapamycin treatment (59, 60) supporting its marker specificity (Fig. 4D). p4E-BP1 level was downregulated in *Mif* mzKO but was rescued by rMif treatment (Fig. 4E). Cd74 KD also inhibited p4E-BP1 level (Fig. 4F). Furthermore, low-dose rapamycin treatment led to decreasing trend in blastocyst size and cell number at both HD and LD culture and blocked the rescue effect by rMif at LD (Fig. S6, D and E). Together, these results suggest that the Mif–Cd74 axis possibly maintains mTORC1 activity to mediate the embryotrophic effects.

## Discussion

Our results identify Mif as an embryo-derived extracellular signal that supports preimplantation development (Fig 4G). Mif accumulated in embryo-conditioned medium, with prominent extracellular Mif detected during the E3.5–E4.5 interval, coinciding with the developmental stage at which loss of Mif produced its strongest phenotype. Genetic depletion of *Mif* or its receptor *Cd74* impaired blastocyst growth and cell proliferation resembling the effects of low-density culture, whereas extracellular Mif activated downstream signaling and reversed the phenotypes. It appears that maternal Mif performs its main function long after the initiation of zygotic control of development (26). Although the relative contributions of regulated and passive release of Mif cannot be completely distinguished by the present experiments, these findings support a model in which extracellular Mif contributes to the signaling environment required for normal blastocyst expansion, rather than serving simply as a marker of embryo abundance. Since Mif was also detected in the human embryo blastocoel fluid (24), it may be important to understand its function in human development and clinical IVF.

We noticed the unchanged Mif protein levels in the *Zp3-Cre;Mif^fl/fl^* and *Gdf9-Cre;Mif^fl/fl^* cKO MII oocytes (Fig. 1D and Fig. S1D). This homeostatic regulation of oocyte Mif levels may be explained by a mechanism where Mif proteins are transported into these oocytes like other maternal proteins (61). Either mechanism may contribute to maintain the maternal Mif protein deposit as a potential mechanism for long-term somatic control of embryonic development.

What may be the developmental advantages of extracellular factors like Mif? Besides its possible contribution to intercellular and inter-embryo communication (4) (Fig. S8), ligand-receptor interactions provide solutions for signal amplification and temporal control. In contrast to the purely reductive division model, early cleavages demand new biomass (62). The demand from the nuclear and plasma membrane compartments both scales exponentially with the round of cell division (*n*) following *N* = *N*_0_ 2*^n^* and *S* = *S*_0_ 2*^n^*^/3^, respectively. Clearly, the demand is at its highest at the blastocyst stage. Concerted release of the sequestered maternal Mif deposit could support the massive cell growth in blastocysts. By sequestering a maternal protein inside cells from its cell-surface receptors, the embryo could control the timing of signaling by regulated release.

The downstream signaling of Mif converges to the PI3K-Akt pathway as many known embryotrophic factors (8, 20). This pathway redundancy may explain the incomplete phenotype caused by *Mif* KO both in vitro and in vivo (Fig. S9). Moreover, the closely related *Ddt/Mif-2* that is capable of binding to Cd74 (28, 29) is also maternally expressed in mouse oocytes and may compensate for the loss of Mif (Fig. S1B). By an unbiased transcriptomic approach, we further connected the downstream signaling of Mif to mTORC1. mTOR is essential for blastocyst expansion and implantation (59, 60, 63–66). Growth regulation by Mif likely goes beyond cell cycle control as *Mif* KO exhibits growth defects despite unchanged EdU^+^ cell percentage (Fig. S2, F and G), possibly involving biomass synthesis through mTORC1 (58). We showed by genetic depletion that Mif and Cd74 were required for maintaining mTORC1 activity at HD culture (Fig. 4, E and F), although mTORC1 pathway upregulation at LD culture was hinted by the negative NES value (Fig. 4C, High density effect, and Fig. S7, C and D) and verified by Western blot of p4E-BP1 levels (Fig. S5D). An unknown mechanism possibly activates mTORC1 to prevent complete developmental arrest at LD culture (Fig. 4G). This was unlikely due to the remaining 21.9 pg ml^-1^ extracellular Mif (Fig. 2B) since acute rMif treatment at 100 pg ml^-1^ failed to activate mTORC1 (Fig. S5C). Future research is warranted to identify this putative compensatory mechanism that may be important for sustaining embryo development.

## Materials and methods

All procedures were approved by the IACUC of Tsinghua University. All embryos were of the C57BL/6J strain and superovulated except for in vivo embryos. Zygotes were cultured at 1 μl^-1^ density in microdrops of KSOM media under mineral oil in 5% CO_2_ in air at 37 °C from E0.5 (Embryonic day 0.5; 24 h post hCG) without changing medium. For low-density culture, embryos were washed three times in KSOM before cultured at or 0.03 μl^-1^ between E3.5–E4.5. Typical setup is 20 embryos in 20 μl (high density) and 1 embryo in 30 μl (low density). See Supplementary Materials for detailed methods.

## Acknowledgments.

We thank Li Yu, Wei Xie, Shunji Jia, Nian Liu, Zhen Jiang, Xukun Lu, Shuyan Ji, Qihan Chen, and Bei Li for valuable critiques; Laboratory Animal Research Center of Tsinghua University for animal care; Hongshuang Li, Fan Lei, Yue Sun, and Jingjing Wang at Imaging Facility for their help. Funding was provided by the National Science Foundation of China 31988101 (to A.M.), Ministry of Science and Technology, National Key R&D Program of China 2023YFA1800301 (to J.X.H.L. and X.W.), and the China Postdoctoral Foundation (to J.X.H.L.).

