## Supporting information for "Oocyte-stored Mif promotes preimplantation development as an embryonic extracellular signal"

### Supporting Information Text

#### Antibodies

Goat anti-Mif (R&D, AF-289-PB); mouse anti-Cd74, LN-2 (SCBT, sc-6262, WB); rat anti-Cd74, IN1 (BD, 555317, IF); rat anti-Cxcr4, 247506 (R&D, MAB21651); mouse anti-Cxcr7/Ackr3, 11G8 (R&D, MAB42273); rat anti-Cd44, IM7 (Invitrogen, 14-0441-81); mouse anti-Cd44/HCAM, DF1485 (SCBT, sc-7297); rabbit anti-Ki67 (Abcam, ab15580, 1:200 for IF); rabbit anti-active caspase-3 (BD, 559565, 1:200 for IF); mouse anti-p21/Cdln1a (SCBT, sc-6246); mouse anti-Cdx2 (Biogenex, MU392A-UC, 1:400 for IF); rabbit anti-Nanog (Cosmobio, RCAB002P-F, 1:200 for IF); rabbit anti-Sox2(Abcam, ab97959); goat anti-Gata6 (R&D, af1700); goat anti-Sox17(R&D, af1924); rat anti-Krt8 (DSHB, TROMA-I); rabbit anti-phospho-Akt S473, D9E (CST, 4060, 1:800 for IF); rabbit anti-Akt, pan, C67E7, (CST, 4691, 1:800 for IF); rabbit anti-phospho-4E-BP1 Thr37/46, 236B4 (CST, 2855, 1:800 for IF, 1:1000 for WB); mouse anti-Rps3 (Proteintech, 66046-1-Ig); rabbit anti-Rpl10 (Proteintech, 17013-1-AP); rabbit anti-Rpl30 (Atlas Antibodies, HPA002651); rabbit anti-Vinculin (Proteintech, 26520-1-AP); rabbit anti-Nlrp5 (Abcam, ab105324); mouse anti-ribosomal RNA (DSHB, y10b); mouse anti-Flag, M2 (Sigma, F1084, 1:1000 for IF). Working concentration is 1 μg $ml^{-1}$ for immunofluorescence and 0.1 μg $ml^{-1}$ for Western blot unless otherwise noted.

#### Inhibitors

LY294002 (Selleck, S1105, working concentration 10 or 25 μM); BX795 (Selleck, S1274, 5 μM); MK2206 (Selleck, S1078, 4 μM); rapamycin (Selleck, S1039, 100 nM); z-VAD-fmk (Selleck, S7023, 20 μM); SB265610 (Selleck, S9516, 400 nM); Plerixafor (TargetMol, T1776, 400 nM); and ACT-1004-1239 (MCE, HY-142617, 4 μM) were dissolved in DMSO as 1000× stock. Prolonged but not acute LY294002 treatment is incompatible with oil overlay and thus was done in 96-well plates (1).

#### siRNAs

siRNA oligo pairs were synthesized with deoxynucleotide overhangs (BGI, Beijing) and injected to zygotes as 10 μM aqueous solution. 5'-3' sequences are as follows.
Negative control,
UUCUCCGAACGUGUCACGUdTdT, ACGUGACACGUUCGGAGAAdTdT (2);
Cd74 siRNA-1 (weaker phenotypes, used throughout this paper),
GGACAUGGAAGACCUAUCUdTdT, AGAUAGGUCUUCCAUGUCCdTdT (3);
Cd74 siRNA-2 (stronger phenotypes),
CCAGGACCAUGUGAUGCAUdTdT, AUGCAUCACAUGGUCCUGGdTdT (4).

#### Mouse genetics

*Mif Cas9-CKO* (*Mif-flox*; T007895) mouse was generated at GemPharmtech, Nanjing on a C57BL/6J background. Germline KO was generated by mating *Mif-flox* with *CMV-Cre* and selecting offspring without the Cre transgene. Germline KO were propogated by either *Mif*$^{+/-}$ ♀ × *Mif*$^{-/-}$ ♂ or *Mif*$^{+/-}$ ♀ × *Mif*$^{+/-}$ ♂ mating. *Zp3-Cre* and *Gdf9-Cre* mice were kindly provided by Wei Xie, Tsinghua University.
Genotyping primer 5'-3' sequences are as follows.
F1, TAGCTGTGGTGCCTCCATTATG;
F2, CAAGGCCCAGCTCAATTTAGG;
R1, ACACGGTCGGATTCTTAACAGC;
R2, CAGTGTGATAGGCAACAGGCTACA.

#### Oocyte and embryo collection

All procedures were approved by the IACUC of Tsinghua University. All mice used were of the C57BL/6J strain. For experiments on WT embryos including proteomics, secretion assay, low-density culture, pharmacological perturbation, and siRNA injection, embryos were fertilized by in vitro fertilization (IVF) to generate large number of embryos. For *Mif* KO phenotypic studies, embryos were fertilized by mating to preserve males for repeated use.
For IVF, virgin female mice (3 weeks weighing 8-12 g) were purchased from Charles River and GemPharmTech. Pregnant mare serum gonadotrophin (PMSG, 5 IU) and human chorionic

gonadotrophin (hCG, 5 IU; Ningbo Sansheng) was injected intraperitoneally 48 h apart. MII oocytes were released from the ampullae 16 h post hCG (hph) into M2 media (Sigma, MR-015-D) containing 0.5 mg $ml^{-1}$ hyaluronidase (Sigma, H4272) covered by paraffin oil (Aibei, M2470). For fertilization, cumulus-oocyte-complexes were released from the ampullae 16 hph into HTF medium (Aibei, M1135) supplemented with 1 mM reduced glutathione (Aibei, M3000) covered by oil and fertilized by sperm capacitated for 1 h in TYH medium (Aibei, M2035); zygotes were washed into KSOM medium (Sigma, MR-121-D or Aibei, M1435) 3 h after adding sperm.
For *Mif* KO oocyte competence test, 3-week-old females were superovulated with 5 IU each of PMSG and hCG and COCs were fertilized similar as above except for skipping glutathione supplementation to HTF medium.
For *Mif* KO in vitro culture, mature females were superovulated with 10 IU each of PMSG and hCG and mated with males. Zygotes were released from ampullae 18 hph.
For *Mif* KO in vivo embryos, mature females were mated with males without superovulation. Blastocysts were released from uterus of plugged females on E3.75 by flushing with M2 medium.

**Culture density**

Zygotes were cultured at 1 $\mu l^{-1}$ (high density) density in microdrops of KSOM media under paraffin oil in 5% $CO_2$ in air at 37 °C from E0.5 (Embryonic day 0.5; 24 h post hCG) to E3.5 without changing medium. Compact morulae were picked out, washed three times in KSOM to remove embryo-derived factors, and cultured at high or low density between E3.5 and E4.5; typical setup is 20 embryos in 20 μl (HD) and 1 embryo in 30 μl (LD, 0.03 $\mu l^{-1}$). The 30 μl significantly dilutes embryo-derived Mif concentration between E3.5 and E4.5, confirmed by ELISA of spent medium.

**Proteomics**

Stable maternal proteome at the two-cell stage was measured separately in the second polar body-associated blastomere (pbB) and the other blastomere (npbB)—a sample collection strategy that aims to facilitate another project. The protein abundance was taken as the mean of that in pbB and npbB. Briefly, zygotes were treated with 100 μg $ml^{-1}$ α-amanitin (Cayman, 17898) from E0.5 to suppress zygotic genome activation. An oil droplet was injected into the zygote near the second polar body thus the two-cell stage blastomere containing the oil droplet was collected as pbB and the other blastomere as npbB between 42 and 46 hph. Zona pellucida was removed by briefly treatment with acidic Tyrode's solution (Sigma, MR-004-D or Leagene, CZ0060); blastomeres were separated by briefly treatment with 0.25% trypsin (Gibco, 25200056 or Yeasen, 40127ES60) and washed three times in KSOM without BSA (Sigma, MR-107). The sample was frozen immediately in liquid nitrogen and stored at -80 °C for downstream processing. Between 135 to 192 blastomeres of each sample were collected as a total of N = 4 pbB and 3 npbB samples (one npbB sample was lost in MS sample preparation).
Label-free mass spectrometry was performed by the Protein Chemistry Facility at the Technology Center of Protein Research, Tsinghua University. Briefly, cells were lysed with 1% sodium deoxycholate in 20 mM Tris pH 8.5, boiled for 10 min at 100 °C, and then sonicated for 30 s in 20 cycles with 30 s interval with rotating at 4 °C. Peptides were digested with porcine trypsin and purified using SDB-RP StageTips. LC-MS/MS were performed on the Q-Exactive HF-X mass spectrometers (Thermo Scientific). The spectrum data were searched against the Uniprot mouse database using the label-free quantification (LFQ) method in the MaxQuant software (5). Relative abundance of each protein was calculated by LFQ value / (sum of LFQ values of all proteins) × 100%.
Secreted proteins were identified from the two-cell maternal proteome by mapping mouse gene symbols to UniProt accessions and integrating UniProt annotations for reviewed status, signal peptide, subcellular localization, transmembrane regions, and lipid/GPI anchoring. For genes with at least one reviewed UniProt accession, only reviewed accession(s) were used for classification and all unreviewed accessions were ignored. Proteins were classified as secreted when reviewed accessions contained a signal peptide (class 1 and 2) and/or explicit extracellular/secreted annotation (class 3). Proteins with secretion evidence together with ER localization (class 4) or transmembrane/lipid-anchor annotation (class 5) were excluded from the soluble secreted set. Genes lacking reviewed accessions were not included in the high-confidence secreted set;

instead, unreviewed secretion annotations were retained as lower-confidence candidates (class 6 and 7).

For re-analysis of zygote proteomics, the average abundance of two samples was taken from the supplementary table in (6).

**Western blot**

Liver extracts were used for antibody validation. Liver was snaped frozen in liquid nitrogen and ground in RIPA buffer. Extract was rotated end to end for 4 h at 4 °C and cleared by centrifugation at 15000 ×g for 10 min. After addition of Laemmli buffer, sample was boiled and run an SDS-PAGE gel. Protein was transferred to nitrocellulose membrane at 300 mA for 1 h in 10% methanol. After blocked in 5% milk in TBST (tris buffered saline with 0.1% Tween-20), membrane was probed by antibodies in 5% milk in TBST and developed by ECL.

Oocytes and embryos were collected directly in 1× Laemmli buffer. Cd74 (LN-2) requires sample denaturation without boiling (1× Laemmli buffer, room temperature for 30 min, otherwise causes protein insolubility) while the sample was denatured at 70°C for 10 min for the other antibodies. Samples were run on a 4-20% gradient Bis-Tris SDS-PAGE gel (LabPAGE, P42015) with MOPS-SDS running buffer. Proteins were transferred to 0.2 μm PVDF membrane, blocked in 5% BSA in TBST, and probed with primary antibody overnight and secondary antibody (donkey IgG H+L HRP conjugates, Proteintech) for 1 h in 5% BSA in TBST, and developed by SuperSignal West Pico PLUS ECL (Thermo, 34577). Membranes were stripped by 2% SDS and 0.8% β-mercaptoethanol (v/v) in 62.5 mM Tris HCl, pH 6.8 at 50°C for 15 min with rotation before re-probing.

**Immunofluorescence**

Embryos were fixed in 4% paraformaldehyde (Dingguo, AR-0211) for 15 min and quenched by 2× vol of TBS. Embryos were transferred to permeabilization buffer (0.1% Triton X-100 and 1% BSA in TBS) and permeabilized for 15 min at room temperature. Morulae and blastocysts were permeabilized with a higher concentration of Triton X-100 (0.25%) for 30 min. All following staining and washing were done in permeabilization buffer. Embryos were incubated with primary antibodies overnight at 4°C, washed twice for 5 min each, incubated with 1 μg $ml^{-1}$ secondary antibodies and DAPI overnight at 4°C, washed once for 5 min. F-actin was stained with 100 ng/mL TRITC-phalloidin (Eaysen, 40734ES75) along with secondary antibodies.

Embryos were washed once and kept in 1% BSA in TBS for 5 min and transferred to oil-overlaid 1 μl drops on a glass-bottom dish (CellVis, D35-20-1.5-N). Embryos were imaged on a Nikon AX R confocal microscope with a 25× silicone oil lens. Z-intensity correction was set up as follows, no correction for 0-15 μm and linear extrapolation beyond 15 μm with 3–4× laser intensity at 85 μm relative to that at 15 μm. The voxel size is typically 0.7×0.7×1 $μm^3$ for blastocyst cell counting and 1.4×1.4×5 $μm^3$ for morphology.

For detergent extraction, oocytes and embryos were incubated in 0.1% Triton X-100 in PBS for 10 minutes, and then fixed by directly adding 1/3 volume of 16% paraformaldehyde (EMS) to the extraction buffer for 15 min.

**ELISA**

For HD, 25 zygotes were placed in 20 μl drops at E0.5 under oil overlay so that embryo count reached around 20 by E3.5. At 1-d interval, the spent drop was collected by pipet and measured for collected volume after removing the embryos. The embryos were washed through 3 drops before transferred to a new drop. Arrested embryos were removed every day and the number of remaining embryos were recorded. Spent medium was cleared by 12000 ×g for 5 min and stored at 4 °C up to 4 d (until all stages were collected). Sample volumes (~20 μl) were brought up to 80 μl with KSOM.

For LD, a single morula was placed in 30 μl drops at E3.5 under oil overlay and spent medium was collected at E4.5. A trial experiment showed that the concentration from individual culture was below the detection limit. Ten drops were thus combined, similarly cleared, and concentrated to 80 μl by 14,000 ×g for 10 min using Amicon Ultra-0.5 (10-kDa cut-off, Sigma, UFC501008).

For MII oocytes, 10 to 25 oocytes were detergent extracted in 10 µl of 0.1% Triton X-100 in PBS, cleared by 12000 ×g for 5 min at 4 °C, and 1–2 µl sample (equivalent of 1–2 oocytes) was used for ELISA measurement.
Mif concentration was measured by a sandwich ELISA assay for mouse Mif per the manufacturer's protocol (Proteintech, KE10027) and normalized by sample volume and embryo count. Briefly, all samples were brought up to 100 µl by the dilution buffer from the kit. The protein standard was dissolved using the same buffer composition as the sample (e.g. KSOM+dilution buffer). The standard curve was fitted with a 4-parameter logistic function of optical density (OD) values for each experiment,

$$OD = LL + \frac{UL-LL}{1+\left(\frac{EC50}{C}\right)^{n}},$$

where *UL*, upper limit; *LL*, lower limit; *EC50*, effective concentration at 50% saturation; *C*, concentration; *n*, Hill coefficient. The standard curve recovery rate was calculated as the ratio between calculated and expected concentrations.

**Mif-mEos live imaging**

Mouse Mif coding sequence was subcloned to the upstream of 3×HA tag, mEos3.1 (7), and auxin induced degron in the pCS2 vector. mRNA in vitro transcribed by mMESSAGE mMACHINE kit (Invitrogen, AM1340) was injected at 150 ng µl$^{-1}$ into zygotes at E0.5. Mif-mEos (green) was photoconverted (magenta) at E1.0 and E3.5 on a FV3000 confocal microscope at 37 °C with 5% $CO_2$ in air, by 405 nm illumination in an embryo-sized ROI at 20-60% laser power and 10 µs pixel$^{-1}$ speed. Converted embryos were immediately tracked by time-lapse imaging at 1-h frame on a Nikon Ti2-E widefield microscope at 37 °C with 5% $CO_2$ in air.

**Recombinant Mif purification**

Full length mouse Mif coding sequence (115 amino acids, N-terminus Met1) was cloned into pET28 plasmid upstream of a HRV3C protease site (LEVLFQ↓GP) and 6× His tag. Rosetta cells (DE3) transformants were grown in Luria-Bertani medium to optical density of 0.6 and induced with 0.5 mM isopropyl β-D-thiogalactoside (IPTG) (Solarbio, I8070) at 16 °C for 4 h. Cell pellet was lysed on ice in 5× pellet volume of lysis buffer (50 mM Tris HCl, pH 8; 300 mM NaCl) with 10 mM imidazole, 1 mg ml$^{-1}$ lysozyme, and 25 U ml$^{-1}$ benzonase (BioSharp, BL1892A) by sonication. Subsequent washing and elution were done with lysis buffer containing various imidazole concentration. Lysate was cleared at 10000 ×g for 30 min. Recombinant proteins were purified under native condition with 1 ml bed volume of Ni-NTA agrose (Qiagen, 30210). Lysate was incubated with agrose at 4 °C for 1 h. After lysate flows through, the resin was washed with 10× bed vol of 10 mM imidazole and 10× bed vol of 20 mM imidazole and eluted with 5× bed vol of 250 mM imidazole. Eluent was sequentially dialyzed for 8 h against 1) 125 mM imidazole and 250 mM NaCl, 2) 50 mM imidazole and 250 mM NaCl, and 3) 150 mM NaCl in 25 mM Tris HCl, pH 7.4 and 5 mM β-mercaptoethanol.
Removing the His-tag was helpful to reduce bacterial contamination. Purified HRV3C protease with 6× His tag was added to purified Mif protein at 1:50 molar ratio and incubated in storage buffer (150 mM NaCl, 25 mM Tris HCl, pH 7.4 and 5 mM β-mercaptoethanol) at 4 °C overnight. After removing protease and uncut Mif protein with Ni-NTA resin, endotoxin was removed by polymyxin B resin treatment (Yeasen, 20518ES10). Briefly, resin was regenerated per manufacturer's protocol and first used to detoxify storage buffer. Then, cut Mif protein was flown through the resin 5 times. Endotoxin level was determined by Limulus Amebocyte Lysate assay (Pierce, A39552S) to be <12.5 EU mg$^{-1}$ protein. Detoxified Mif protein was concentrated to 1 mg ml$^{-1}$ in storage buffer and snap frozen in aliquots. After thawing, the protein was cleared by 12000 ×g for 5 min at 4 °C and its concentration was remeasured before use.
As a control, rMif was boiled for 5 min at 95 °C and cleared by 12000 ×g for 5 min.

**Recombinant Mif treatment**

Acute treatment was routinely done by washing embryos in KSOM three times and then at 100 pg ml$^{-1}$ rMif in KSOM for 30 min. This condition was sufficient to initiate downstream signaling even in embryos at high density (Fig. 3 and 4). In contrast, chronic treatment of WT embryos at high

density E0.5–4.5 had minimal effects on development possibly due to embryo-derived Mif (Fig. 2).
Rescue experiment at LD culture was done by washing embryos in KSOM three times and then in 100 pg $ml^{-1}$ rMif in KSOM three times before culturing in the latter medium.
The concentration for rescue experiment E0.5–4.5 is determined as follows. At HD culture (1 $\mu l^{-1}$), 2.1 pg released Mif per embryo would give a concentration of 2.1 ng $ml^{-1}$. Given that recombinant Mif is three-fold less potent than endogenous Mif (8), 6.3 ng $ml^{-1}$ of rMif would function equivalently to the endogenous Mif after factoring in a three-fold excess.

**EdU assay**
At the end of rMif treatment, embryos were washed by KSOM, treated with 5-ethynyl-2'-deoxyuridine (EdU, TargetMol, T17341) at 50 µM in KSOM for 1 h, and fixed directly in PFA. Recombinant Fgf2 (Preprotech, AF-100-18B, 40 µg $ml^{-1}$) with heparin (1 µg $ml^{-1}$) was used as positive control. Permeabilization was done as in immunofluorescence. Click chemistry conjugation to azide-AF565 was done with Cell-Light EdU Apollo In Vitro Kit (RiboBio, R11053.12) for 30 min before performing immunofluorescence (9).

**TUNEL assay**
PFA-fixed embryos were labeled by RiboAPO TUNEL Apoptosis Detection Kit (RiboBio, C11026) before performing immunofluorescence. Permeabilization was done as in immunofluorescence. DNAse I treatment on fixed embryos at 0.1 U $\mu l^{-1}$ for 3 h was used as positive control.

**Developmental rate**
Oocyte competence was calculated as two-cell embryo / COC count.
Embryonic developmental progress was examined from E1.5 at 1-d interval. Developmental rate was normalized to two-cell embryo stage. To avoid zero count and allow robust statistical test (Fisher's exact test), data were collapsed to 2×2 contingency table of normal and arrested embryo counts in each experimental condition. The normal criteria are as follows: two-cell stage at E1.5; more than 4 cells at E2.5; morulae and blastocysts at E3.5; blastocysts at E4.5. Degenerated and dead refer to embryos with dead blastomeres; fragmented refers to embryos with live but uncompact blastomeres.

**PI(3,4,5)$P_3$ sensor**
Mouse Grp1 PH domain (aa 2-154) was subcloned downstream of MAPKK NES sequence (MALQKKLEELELD) and mScarlet-I in the pXT7 vector. mRNA in vitro transcribed by mMESSAGE mMACHINE kit (Invitrogen, AM1344) was injected at 180 ng $\mu l^{-1}$ to zygotes (1). CAAX-GFP mRNA was co-injected for visualizing junctional colocalization but omitted in Grp1 PH probe intensity measurement experiments as CAAX-GFP interferes with Grp1 PH probe localization. Embryos were washed in KSOM three times and imaged in 10 µl KSOM drops under oil at 37 °C with 5% $CO_2$ in air on a FV3000 confocal microscope with a 40X air lens.

**Blastocyst outgrowth**
E4 blastocysts were individually placed in 1 ml medium in glass bottom dishes coated with 0.2% gelatin for up to 4 d. The medium contains 15% fetal bovine serum, 1000 IU $ml^{-1}$ recombinant LIF, 0.1 mM non-essential amino acids, 2 mM L-glutamine, 0.1 mM β-mercaptoethanol, and 1× penicillin/streptomycin.

**RNA-seq and data analysis**
We collected samples at E4.0 or 12 h after switching to LD and/or supplementing rMif, because this duration should be adequate for transcriptional responses. Although the failure in morula-to blastocyst transition is a phenotype of Mif mzKO and Cd74 KD, mixing blastocysts with arrested morulae would lead to transcriptional changes dominated by stage differences (Fig. S5D). Thus, we only collected blastocysts.
We built bulk cDNA library in house by Poly(A) based reverse transcription, reverse transcriptase based tagmentation, and PCR based amplification. Briefly, 10 blastocysts were removed of zona by Tyrode's solution, washed 3 times in nuclease-free PBS + 0.3% polyvinylpyrrolidone, and

collected in the lysis buffer. Full-length cDNA library was made per the manufacturer's protocol with an PCR amplification cycle number of 14 (Single Cell Full Length mRNA-Amplification Kit, Vazyme, N712). Half ng of cDNA was used for tagmentation per the manufacturer's protocol with an PCR amplification cycle number of 13 (TruePrep DNA Library Prep Kit V2 for Illumina, Vazyme, TD503). Fragments containing ~330 bp insert length were purified and sent for Hiseq-PE150 Illumina sequencing at Novogene (Beijing). Each sample was sequenced for 1 GB data. We analyzed data as follows. We performed quality control and adapter trimming using Trim Galore (v0.6.10) with a Phred quality threshold of 20, minimum read length of 36 bp, and stringency of 3 bp for adapter matching. Trimmed reads were aligned to the mouse GRCm38 reference genome (GENCODE vM23 annotation) using STAR (v2.7.11a) with 12 threads, allowing up to 20 multimapping hits, a maximum mismatch rate of 4% per read, and intron size range of 20 bp to 1 Mb. Gene level counts were quantified from the aligned BAM files using featureCounts (v2.0.6). After removing lowly expressed genes (≥2 samples with count ≥ 5), differential expression of protein coding genes was modeled by condition or by condition + batch (for E4.0 Mif WT and mzKO) using Deseq2 (v1.38.3). The following batches were modeled separately with the first condition as reference: siCtrl, siCd74, Boiled, and rMif (E4.0); Mif WT and mzKO (E4.0). Principal component analysis (PCA) was performed using DESeq2 variance-stabilized counts (VST). Differentially expressed genes (DEG) were filtered by the absolute Log2 fold change (LFC), |LFC| ≥ 1, and adjusted $p < 0.05$, which was also used for volcano plot. Raw Wald's statistic from DESeq2 was used for GSEA analysis by clusterProfiler (v4.15.0.3) against the Molecular Signatures Database (MSigDB) (v7.5.1); Hallmark Pathway gene sets (N = 50) were screened by adjusted $p < 0.05$ (Benjamini-Hochberg correction). GO term analysis was performed using clusterProfiler against the mouse genome-wide annotation database, org.Mm.eg.db (v3.16.0). For TPM calculation, low-expression genes expressed in <50% of samples in each condition were removed. Counts were normalized to TPM and used for heatmap comparing samples.

**Image analysis**

*Mif IF intensity across stages.* Embryos across stages were processed and imaged in the same experiment. Embryos were segmented in Imaris using the Surface function to calculate mean intensity. WT and mKO zygotes were set to 1 and 0, respectively, and other stages were normalized to them.

*Mif-mEos intensity.* Embryos were tracked by the unconverted channel in ImageJ using the TrackMate plugin to calculate mean intensity (10). Only embryos that were trackable throughout the movie were kept for analysis. Note that mainly the thicker ICM region was detected in blastocysts due to the lower intensity in the thinner TE region in widefield images, thus the region heterogeneity may not be fully accounted for by this plugin.

*Blastocyst size.* Blastocyst cross-sectional area is measured by manually fitting the embryo with one or more ovals in ImageJ.

*PI(3,4,5)P$_3$ sensor intensity.* Junctions close to the morula surface on the lens (brighter) side of the morula were measured, excluding cavitating embryos. Segmented lines were manually drawn along the junction and mean intensity of all pixels were taken.

*pAkt intensity.* Similar to PI(3,4,5)P$_3$ sensor but the line was drawn in the phalloidin channel to measure pAkt channel.

*p4E-BP1 and Akt intensity.* Embryos were segmented in Imaris using the Surface function by the phalloidin channel when available to calculate mean intensity of the whole embryo excluding the cavity region.

*Blastocyst cell number.* DAPI channel was segmented using the MINS software (11). Other marker identities were determined by their respective intensity threshold set manually and consistently across experimental conditions.

*Temporal analysis of cell number.* The intensity threshold of cell fate markers Cdx2 and Sox2 were determined separately for each stage. Overall, their intensity was low at E3.5. The increase of Cdx2$^+$ and Sox2$^+$ cell numbers are likely contributed by both cell proliferation and cell fate inclusion due to either upregulation of marker expression or real cell differentiation events. The decrease in Cdx2$^-$Sox2$^-$Sox17$^-$ cell number is likely contributed by unspecified cells acquiring fate markers in control and cell proliferation of unspecified cells or cells losing fate markers in Cd74 knockdown.

*Cell status coordination.* Although we did not observe clear separation between G1 and G2/M phase on the EdU–DAPI scatter plot (12) , this plot is still useful to show cell status relative to DNA synthesis/S phase. We calculate Euclidean distance between cell pairs on the plot to compare cell status coordination.

GPT-6 and DeepSeek were used to help write R, ImageJ, and MATLAB code for image analysis and data plotting.

**Statistics**

For single comparison, two-sided Welch's t-test, which tolerates non-normal distribution, was used; except for AM/WT ratio, two-sided Student's t-test was used for paired t-test. The choice of test for multiple comparisons was chosen based on the published guidelines (13). Dunnett's test was used for multiple comparisons with a single control. Two-sided Welch's t-test with Holm correction was used for multiple comparisons with different controls. P-value stars: $p \leq 10^{-4}$, ****; $p \leq 10^{-3}$, ***; $p \leq 10^{-2}$, **; $p \leq 0.05$, *; $p > 0.05$, ns, not significant. All boxplots show median and interquartile range (IQR) with whiskers marking the inlier range.

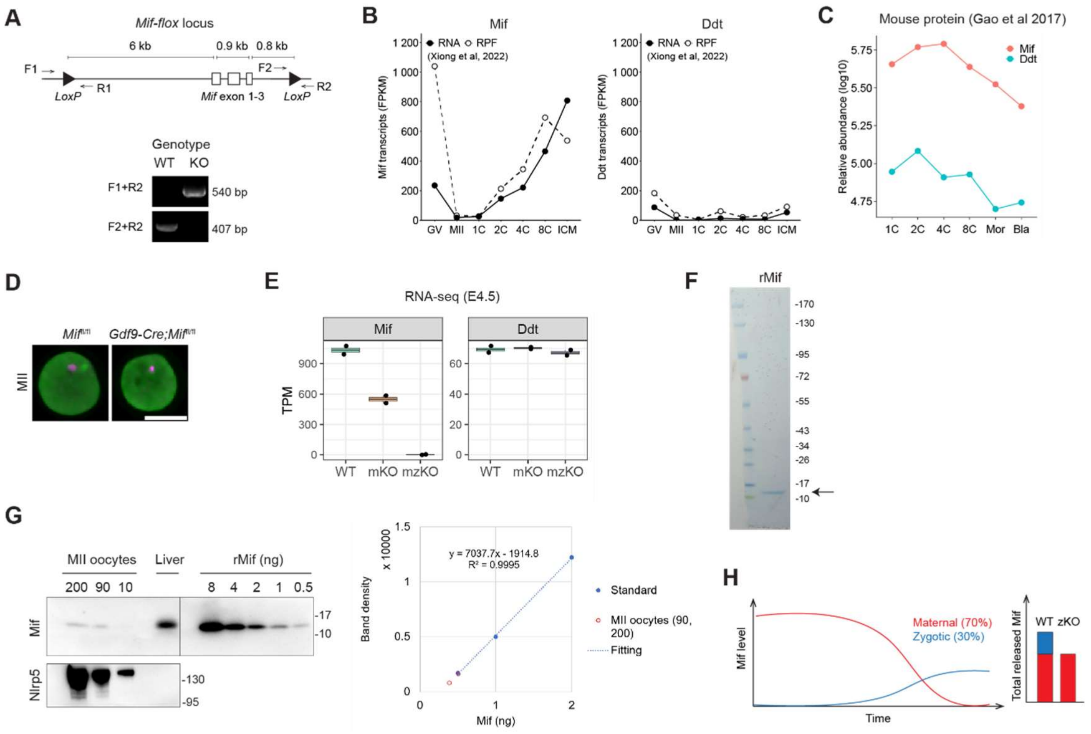


**Figure S1.** Expression and knockout of Mif.

**A:** Genomic locus of *Mif*$^{flox}$ allele and genotyping results of wildtype and *Mif*$^{-/-}$ tail clips by PCR. Note that Cre excises the whole gene.

**B:** mRNA and ribosome-protected fragment (RPF) level Mif receptors according to published data (14). GV is fully grown oocytes; 1C is at PN5 stage; 2C is at late 2C; 4C is around E1.75; 8C is around E2.0; inner cell mass (ICM) of blastocyst is around E3.5.

**C:** Relative protein abundance of mouse Mif and Ddt according to published data (6). Note that Mif and Ddt levels decrease as development progresses.

**D:** Immunofluorescence of Mif (green) and DAPI (magenta). *Mif*$^{fl/fl}$ and *Gdf9-Cre;Mif*$^{fl/fl}$ MII oocytes. Note that protein level remains unchanged. Bar, 50 μm.

**E:** Mif and Ddt mRNA levels from RNA-seq of E4.5 *Mif* WT, mKO, and mzKO blastocysts. Note that Mif mRNA is absent in mzKO; mKO that lacks one allele expresses half of the Mif mRNA in WT suggesting that there is little feedback regulation on Mif transcription due to KO; Ddt level remains unchanged in mKO and mzKO.

**F:** Coomassie stain of purified recombinant Mif (rMif) shows protein purity.

**G:** Western blot of 200, 90, and 10 MII oocytes, liver extract, and rMif protein standard. Nlrp5, loading control. By extrapolating the plot for 200 and 90 oocytes, each oocyte is estimated to contain 2.5 and 4.3 pg Mif, respectively.

**H:** Cartoon of maternal and zygotic contribution of Mif.

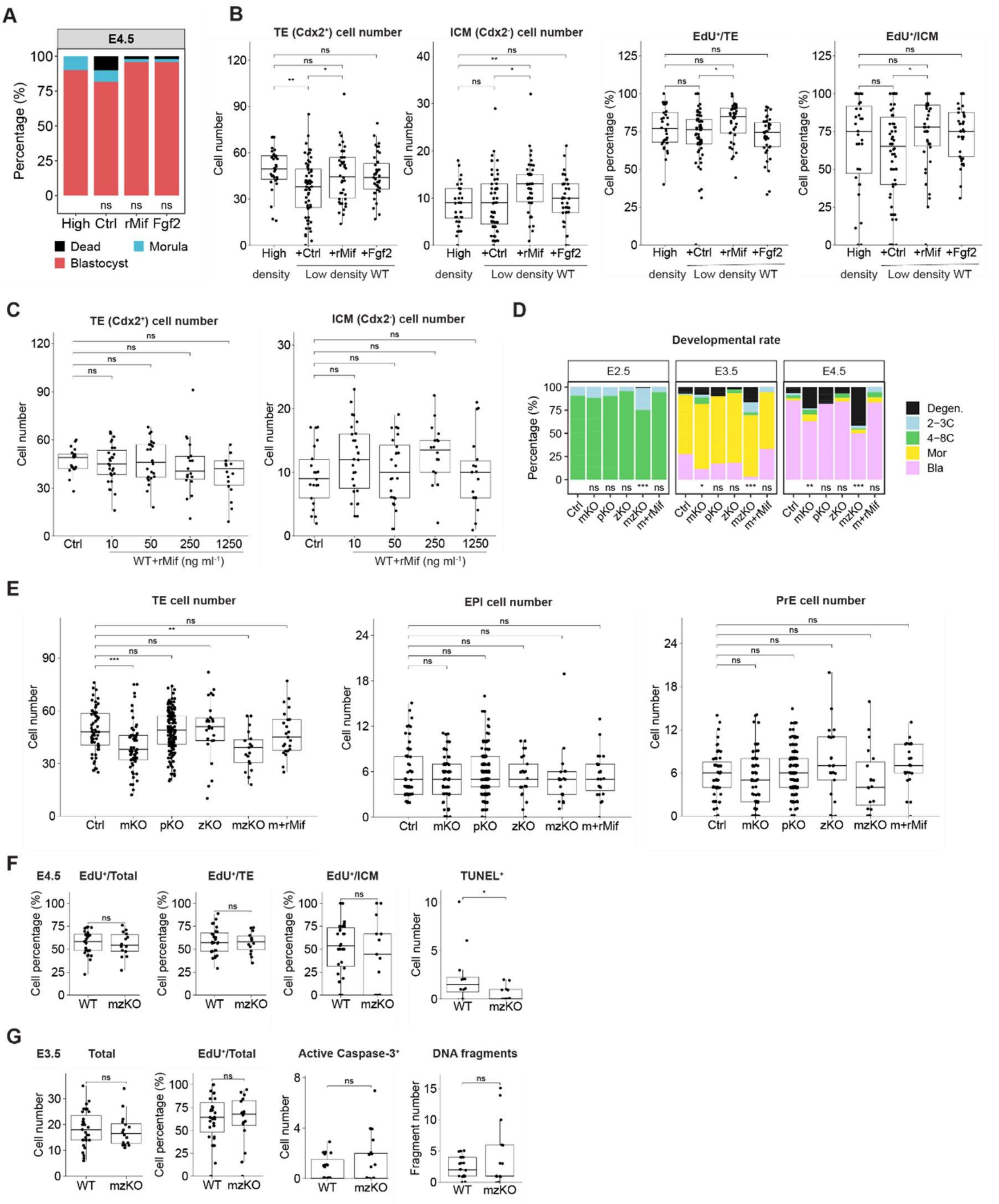


**Figure S2.** In vitro phenotypes of *Mif* knockout.

**A:** (A and B) WT morulae were cultured at the high or low density in the presence of boiled (Ctrl) or normal rMif E3.5-4.5. Fgf2 is a positive control. Developmental rate normalized to morula stage; N = 20, 49, 48, and 48; Fisher's exact test. Note that low density slightly decreased blastocyst formation rate.

**B:** Cell number and EdU incorporation by lineage in E4.5 blastocysts. TE, trophectoderm; ICM, inner cell mass. N = 32, 59, 42, and 39; two-sided Welch's t-test with Holm correction. Note that low density decreased TE cell number but was rescued by rMif.

**C:** WT zygotes at high density were treated with 250 ng $ml^{-1}$ boiled rMif (Ctrl) or indicated concentration of normal rMif E0.5–4.5. E4.5 blastocyst cell number; N = 21, 27, 24, 18, and 18; Dunnett's test. Note that rMif did not change TE or ICM cell number at high embryo density.

**D:** *Mif* KO zygotes were cultured at high density E0.5–4.5. Developmental rate normalized to the two-cell stage. Ctrl (*Mif*$^{+/-}$ ♀ × WT ♂); mKO (maternal KO; *Mif*$^{-/-}$ ♀ × WT ♂); pKO (paternal KO; WT ♀ × *Mif*$^{-/-}$ ♂); zKO (zygotic KO; *Zp3-Cre;Mif*$^{fl/fl}$ ♀ × *Mif*$^{-/-}$ ♂); mzKO (maternal-zygotic KO; *Mif*$^{-/-}$ ♀ × *Mif*$^{-/-}$ ♂); m+rMif (mKO treated with 6.3 ng $ml^{-1}$ recombinant Mif E0.5–4.5). Degen., degenerated; mor, morula; bla, blastocyst. N = 90, 57, 155, 44, 172, and 18; Fisher's exact test. Note that mKO and mzKO inhibited blastocyst formation even at high density.

**E:** Cell number by lineage in E4.5 *Mif* KO blastocysts at high density. Cdx2$^{+}$ (TE), Sox2$^{+}$Sox17$^{-}$ (EPI, epiblast), and Sox17$^{+}$ (PrE, primitive endoderm). N = 59, 63, 137, 25, 19, and 23; Dunnett's test. Note that mKO and mzKO decreased TE cell number even at high density.

**F:** EdU$^{+}$ (cell proliferation marker) cell percentage (N = 27 and 13) and TUNEL$^{+}$ (apoptotic marker) cell number (N = 12 and 18) in E4.5 *Mif* KO blastocysts; two-sided Welch's t-test. Note that mzKO did not change EdU incorporation.

**G:** Total cell number and EdU$^{+}$ cell percentage (N = 31 and 16); active caspase-3$^{+}$ (apoptotic marker) cell number and DNA fragment count (N = 23 and 19) in E3.5 *Mif* KO embryos; two-sided Welch's t-test. Note that mzKO did not change EdU incorporation.

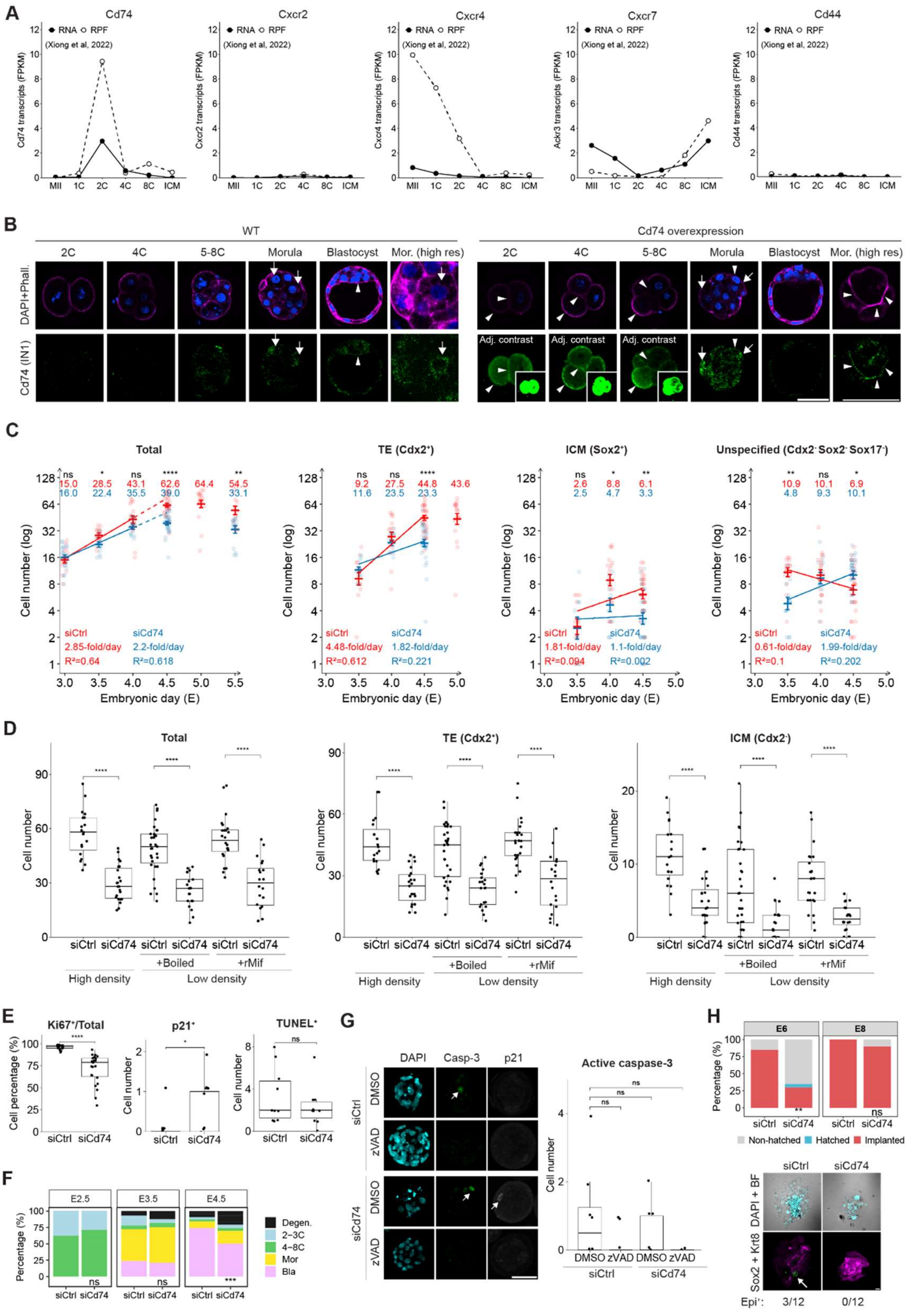


**Figure S3.** Expression and knockdown of Cd74.

**A:** mRNA and ribosome-protected fragment (RPF) level Mif receptors according to published data

(14). 1C is at PN5 stage; 2C is at late 2C; 4C is around E1.75; 8C is around E2.0; inner cell mass (ICM) of blastocyst is around E3.5. Note that Cd74 and Cxcr7 are expressed zygotically and Cxcr4 maternally.

**B:** Immunofluorescence of Cd74 (clone IN1) (green), phalloidin (magenta), and DAPI (blue) in MII oocytes and preimplantation embryos. Sample collection timing is the same as in Fig.1; 4C and 6-8C are both E2.5. Cd74 mRNA were injected to zygotes for overexpression (OE). Arrow and arrowhead, Cd74 in the perinuclear/nuclear and plasma membrane regions, respectively. High resolution images of WT and OE show perinuclear/nuclear region and apical junction in morula, respectively. 2C to 8C OE images were adjusted for contrast to avoid saturation, with the original images shown in the inset. Note the junctional localization in OE morulae.

**C:** Temporal dynamics of cell number in log scale. The horizontal line and error bar show mean ± s.e.m.; N = 20; two-sided Welch's t-test with Holm correction between siCtrl (red) and siCd74 (blue). Solid line is linear regression of cell number; dotted line is extrapolated from solid line to show prediction. Note that the cell number in siCd74 could not catch up when cultured beyond E4.5. The intensity threshold of cell fate markers Cdx2 and Sox2 were determined separately for each stage. Overall, their intensity was low at E3.5. The increase of $Cdx2^{+}$ and $Sox2^{+}$ cell numbers are likely contributed by both cell proliferation and cell fate acquisition events. The decrease in $Cdx2^{-}Sox2^{-}Sox17^{-}$ cell number is likely contributed by unspecified cells acquiring fate markers in control and cell proliferation of unspecified cells or cells losing fate markers in Cd74 knockdown.

**D:** Morulae were treated at high or low density in the presence of 100 pg $ml^{-1}$ boiled (Ctrl) or normal rMif E3.5-4.5. The plots show cell number in blastocysts; N = 19, 23, 31, 21, 24 and 20; two-sided Welch's t-test with Holm correction. Note that Cd74 knockdown partially blocked the rescue of cell number by rMif in the low-density culture, especially in ICM.

**E:** Cell proliferation (Ki67), p21, and apoptosis (TUNEL) analysis in E4.5 blastocysts at high density. N = 19 and 23 (Ki67); 8 and 8 (p21); 10 and 10 (TUNEL); two-sided Welch's t-test. Note that Cd74 KD inhibited cell proliferation.

**F:** Developmental rate normalized to the two-cell stage. N = 72 and 81; Fisher's exact test. Note that Cd74 KD inhibited blastocyst formation.

**G:** Immunofluorescence of DAPI (cyan), active caspase-3 (green) and p21 (white) in embryos treated with apoptosis inhibitor z-VAD-fmk (15) E3.5-4.5. The plot shows apoptotic cell number; N = 8, 9, 8, and 5; Dunnet's test. Note that zVAD suppressed apoptosis in both siCtrl and siCd74 validating the specificity of active caspase-3 staining.

**H:** E4 blastocysts were cultured with serum and Lif. The bar plot shows the morphology at E6 and E8; N = 20 and 20; Fisher's exact test. Note that Cd74 KD delayed implantation. The images show immunofluorescence of Sox2 (green, epiblast marker, arrow), and Krt8 (magenta, trophoblast marker) in E8 implanted outgrowth. Arrow, epiblast. The percentage of outgrowth with epiblast is 25% in siCtrl and 0 in siCd74; all outgrowth formed trophoblast. All experiments in this figure were done with siRNA-1. Scale bar, 50 μm.

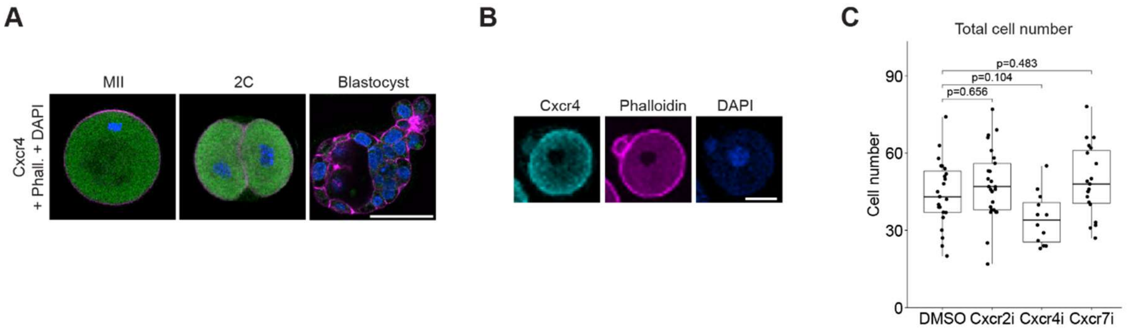


**Figure S4.** Expression and perturbation of Mif co-receptor Cxcr4.

**A:** Immunofluorescence of Cxcr4 (green), phalloidin (magenta), and DAPI (blue) in MII oocytes and preimplantation embryos.

**B:** Cxcr4-Flag mRNA was injected to zygotes. Immunofluorescence of Flag-tag (cyan), phalloidin (magenta), and DAPI (blue) in zygotes.

**C:** Embryos were treated with 400 nM SB265610 (Cxcr2 inhibitor), 400 nM Plerixafor or (AMD3100; Cxcr4 inhibitor), and 4 µM ACT-1004-1239 (Ackr3/Cxcr7 inhibitor) E0.5-4.5. The plot shows total cell number in E4.5 blastocysts; N = 23, 25, 12, and 19; Dunnet’s test. Bar, 50 µm.

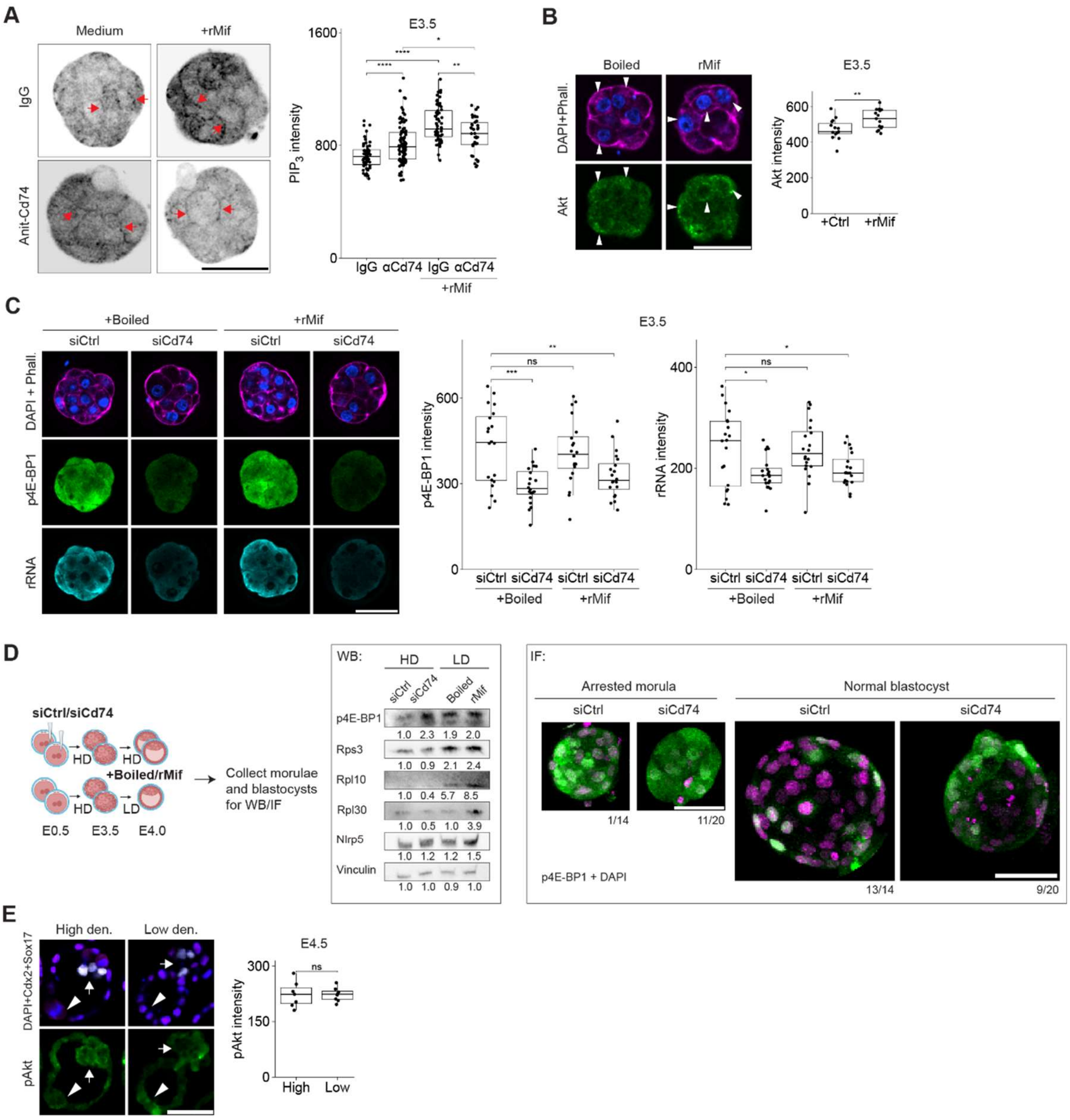


**Figure S5.** PI3K-Akt-mTORC1 pathway activity.

**A:** Zygotes were injected with mScarlet-PH(Grp1) (PI(3,4,5)$P_3$ probe) mRNA. PI(3,4,5)$P_3$ is enriched at cell-cell junctions in morulae (arrows) shown by the inverted grayscale images. Morulae were treated with 100 µg ml$^{-1}$ IgG control or anti-Cd74 neutralizing antibody (clone LN-2) that recognizes its extracellular domain (16) for 30 min, and then stimulated with 100 pg ml$^{-1}$ rMif in the presence of antibodies for 5 min. The plot shows junction intensity; N = 57, 92, 64, and 38; two-sided Welch's t-test with Holm correction. Note that rMif treatment at the physiological concentration strongly upregulated junctional PI(3,4,5)$P_3$ intensity, which was partially blocked by the neutralizing antibody, directly supporting the extracellular mode of action of Mif and linking Cd74 at its cell-surface receptor. Although inhibitory at high concentrations (50–100 μg ml$^{-1}$), LN-2 has also been used to activates the PI3K pathway at lower concentrations (17). The junctional localization of Cd74 may have limited the antibody access to the receptor due to tight junctions, effective lowering the local antibody concentration and allowing it to function as a partial agonist. Nonetheless, PI(3,4,5)$P_3$ activation by rMif in the presence of LN-2 was lower than with control IgG.

**B:** E3.5 morulae treated with 100 pg ml$^{-1}$ boiled or normal rMif for 30 min. Immunofluorescence of DAPI (blue), phalloidin (magenta), and Akt (green). The plot shows whole-embryo intensity; N = 15 and 15; two-sided Welch's t-test. Note that rMif promotes Akt localization at perinuclear regions (arrowhead), unlike pAkt which is enriched at cell-cell junction (Fig. 3).

**C:** Immunofluorescence of DAPI (blue), phalloidin (magenta), phospho-4E-BP1 T37/46 (green), and ribosomal RNA (cyan) in E3.5 morulae treated with 100 pg ml$^{-1}$ boiled or normal rMif for 30 min. The plot shows whole-embryo intensity; N = 20, 20, 20 and 20; Dunnett's test. Note that acute rMif treatment at this concentration did not activate mTORC1 activity; Cd74 KD decreased ribosomal RNA level.

**D:** siCtrl and siCd74 were cultured at high density while WT were at low density and supplemented with 100 pg ml$^{-1}$ boiled or normal rMif E3.5–4.0. Both blastocysts and arrested morulae were collected. Western blot (113 embryos per condition) and immunofluorescence were performed on the same batch of embryos. In WB, Nrlp5 and vinculin are loading control; note that LD activated mTORC1 activity and ribosomal protein production (Boiled versus siCtrl) which were further enhanced by rMif supplementation (rMif versus Boiled); siCd74 also showed overall higher p4E-BP1 level compared to siCtrl, which is explained by the IF results. In IF of DAPI (magenta) and p4E-BP1 (green), p4E-BP1 was downregulated in blastocysts compared to morulae; siCd74 arrested more embryos at morula stage causing overall higher p4E-BP1 level. This observation supports the rationale of collecting only blastocysts for RNA-seq to prevent cellular states being dominated by stage differences (Fig. 4).

**E:** Immunofluorescence of DAPI (white), Cdx2 (white), Sox17 (magenta), and pAkt (green) in embryos cultured at high or low density E3.5–4.5. The plot shows whole-embryo pAkt intensity; N = 7 and 8; two-sided Welch's t-test. Arrow, ICM; arrowhead, TE. Note that Akt was still activated at low density. Scale bar, 50 μm.

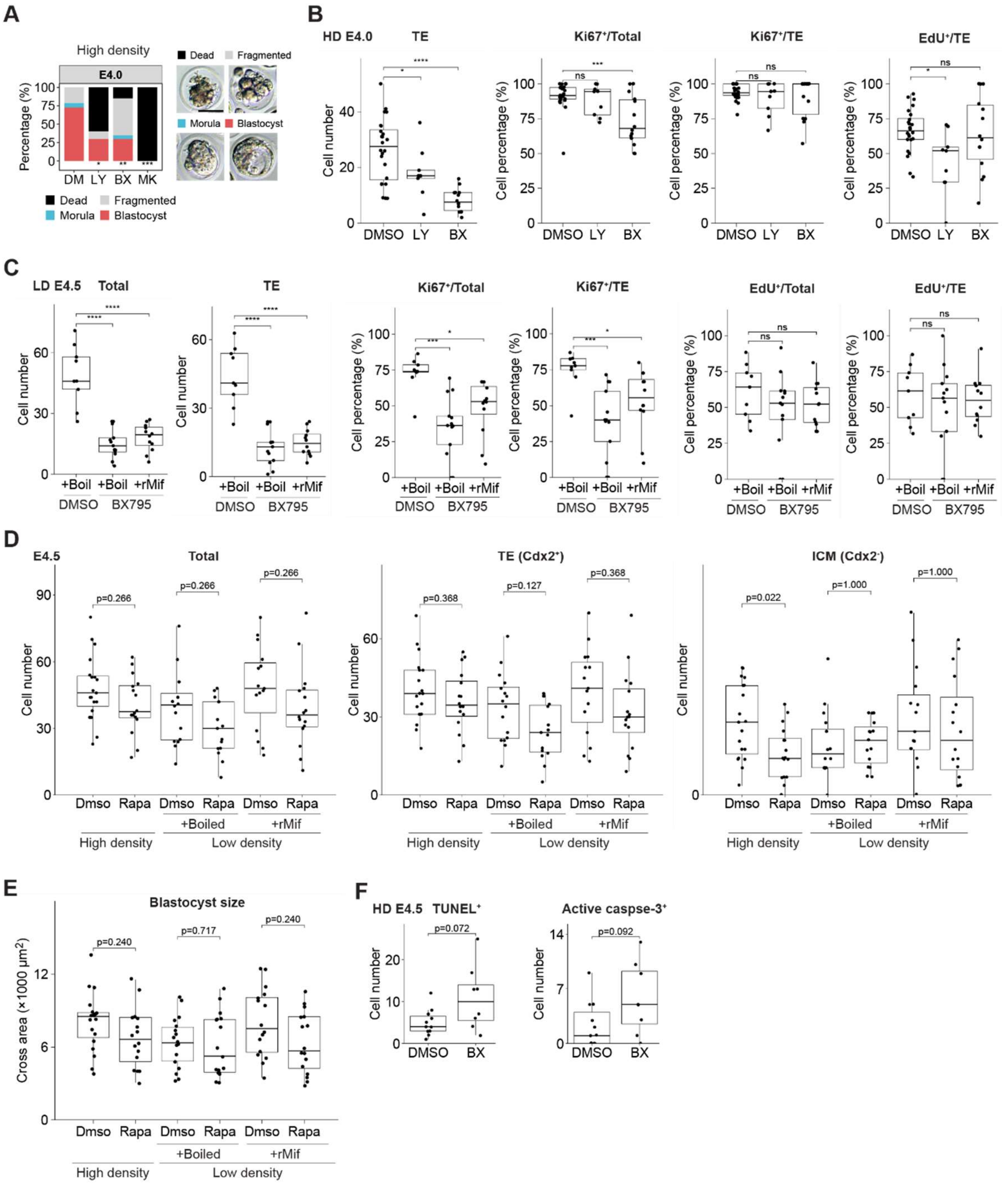


**Figure S6.** Pharmacological perturbations of PI3K-Akt-mTOR pathway.

**A:** (A and B) Morulae were treated with DMSO, 10 μM LY294002 (LY, PI3K inhibitor), 5 μM BX795 (BX, PDK1 inhibitor), or 4 μM MK2206 (MK, Akt inhibitor) at high density E3.5–4.0. The plot shows developmental rate; N = 33, 10, 20, and 20; Fisher's exact test. Note that MK killed all morulae while LY and BX partially inhibited blastocyst formation.

**B:** Cell number and percentage. N = 21, 5, and 13; Dunnett's test. See also Fig. 3. Note that LY and BX inhibited blastocyst growth even at high embryo density suggesting that PI3K and PDK1 are required for blastocyst growth.

**C:** WT morulae were treated with DMSO or 5 μM BX795 in the presence of 100 pg $ml^{-1}$ boiled or normal rMif at the low density E3.5–4.5. N = 9, 13, and 12; Dunnett's test. Note that BX bloecked Mif's rescue of blastocyst growth at low embryo density suggesting PDK1 is downstream of Mif.

**D:** (D and E) Morulae were treated with 100 nM rapamycin in the presence of 100 pg $ml^{-1}$ boiled or normal rMif at high or low density E3.5–4.5. N = 19, 16, 16, 15, 15, and 16; two-sided Welch's t-test with Holm correction. Note that Rapa slightly inhibited blastocyst growth even at high density and blocked Mif's rescue of blastocyst growth at low embryo density suggesting mTORC1 is required for blastocyst growth downstream of Mif. It also means that mTORC1 is likely still active at LD culture.

**E:** Cross-sectional area. N = 19, 16, 17, 17, 16, and 16; two-sided Welch's t-test with Holm correction.

**F:** WT morulae were treated with DMSO or 5 μM BX795 at high density E3.5–4.5. N = 11 and 8; two-sided Welch's t-test.

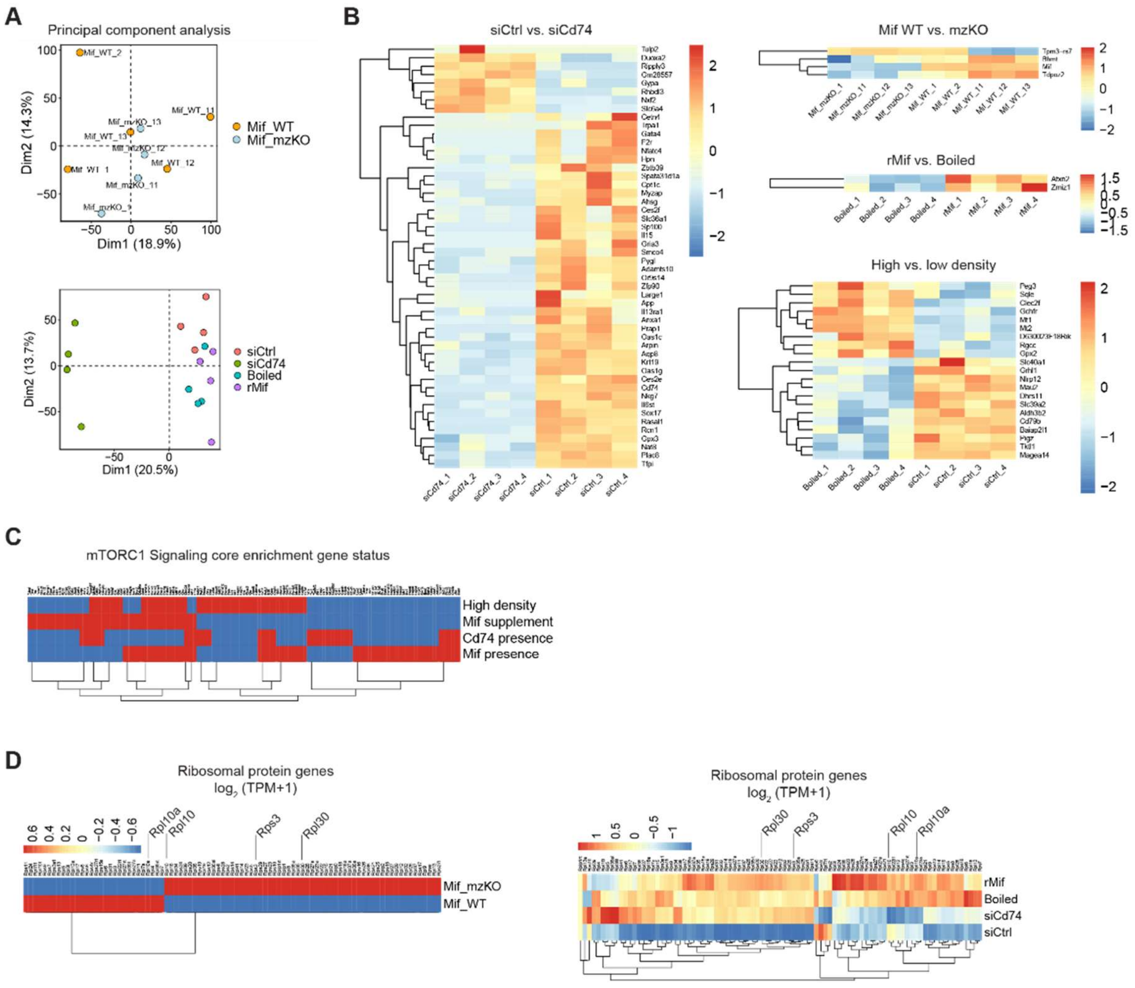


**Figure S7.** Transcriptomics of *Mif* KO, *Cd74* KD, and low-density culture.

**A:** PCA. Mif 1-3 and 11-13 were from two batches; the bottom plot was from another batch.

**B:** Heatmaps of top 50 DEG expression in $\log_2$ (TPM+1).

**C:** The core enrichment genes in each comparison are in red while other genes in blue. Note that the 141 overlapped genes were shown (gene set = 201 genes); the four comparisons had only 1 common core enrichment gene, *Dhcr24*. “Mif presence” and “Cd74 presence” drove the upregulation of a subset of mTORC1 pathway genes while “Mif supplementation” and “High density” drove the downregulation of a different subset.

**D:** Expression level of ribosomal protein (RP) genes; column-normalized Z-score of each gene. N = 84 genes. Labeled genes were verified by Western blot. Note that LD culture transcriptionally upregulated 98% of RP genes, with three of them verified by Western blot (Fig. S5D, Boiled versus siCtrl); Mif supplementation at LD further upregulated 64% of RP genes (Fig. S5D, rMif versus Boiled). mTORC1 possibly promotes ribosome biogenesis at LD culture (18).

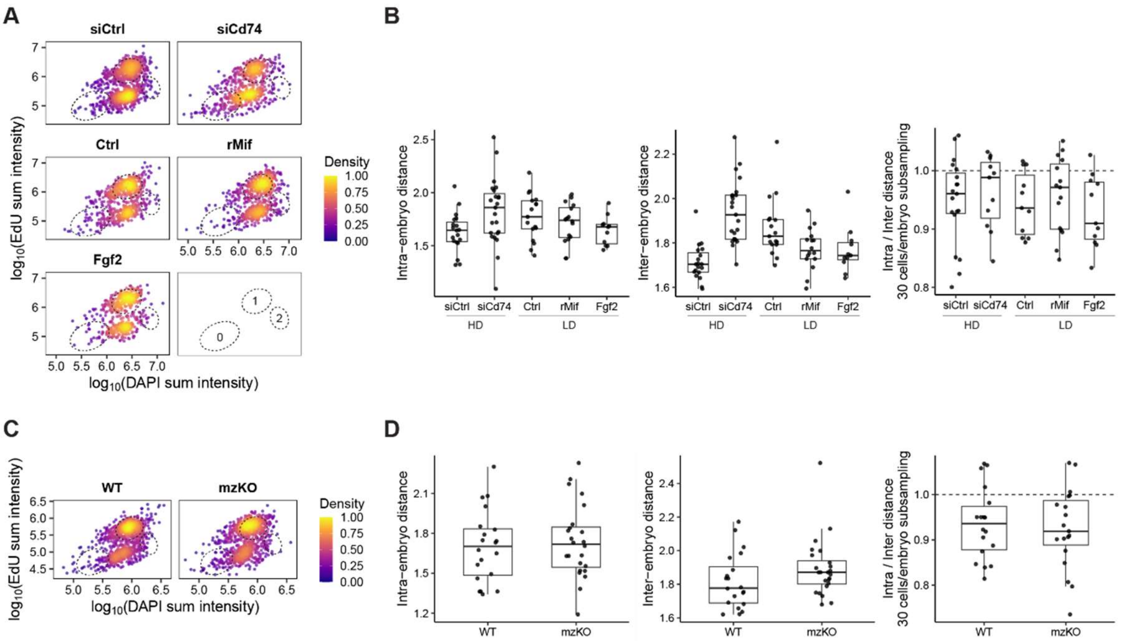


**Figure S8.** Intra- and inter-embryo coordination of cell status.

**A:** Status of each cell's EdU versus DAPI intensity from confocal images in E4.5 blastocysts. siCtrl and siCd74 were at high density; WT morulae were cultured individually from E3.5 in 30 µl drops containing 100 pg $ml^{-1}$ boiled (Ctrl) or normal rMif, and Fgf2 (40 µg $ml^{-1}$, with 1 µg $ml^{-1}$ heparin). Dotted circles, non-replicative (0); replicating (1); replicated (2) cells. Scale, relative density. N = 1091, 685, 648, 737, and 510 cells (pooled from 19, 23, 17, 16, and 11 blastocysts). Note the heterogeneity was higher in Cd74 KD and low density where cells shifted to circle "0", which was rescued by rMif and Fgf2.

**B:** Mean Euclidean distance between any two cells on the EdU-DAPI plot. From left, distance between cells within (intra) and between (inter) embryos and each embryo's intra and inter-embryo distance ratio. Note that cell status in Cd74 KD and low density was more heterogenous, both within and between embryos, which was partially rescued by rMif and Fgf2.

**C:** Same as in (A) but for *Mif* KO E4.5 blastocysts at high density. N = 1061 and 1038 cells (pooled from 20 and 24 blastocysts). Note the higher heterogeneity in mzKO where cells shifted to circle "0".

**D:** Same as in (B). Note that cell status between mzKO embryos was more heterogenous.

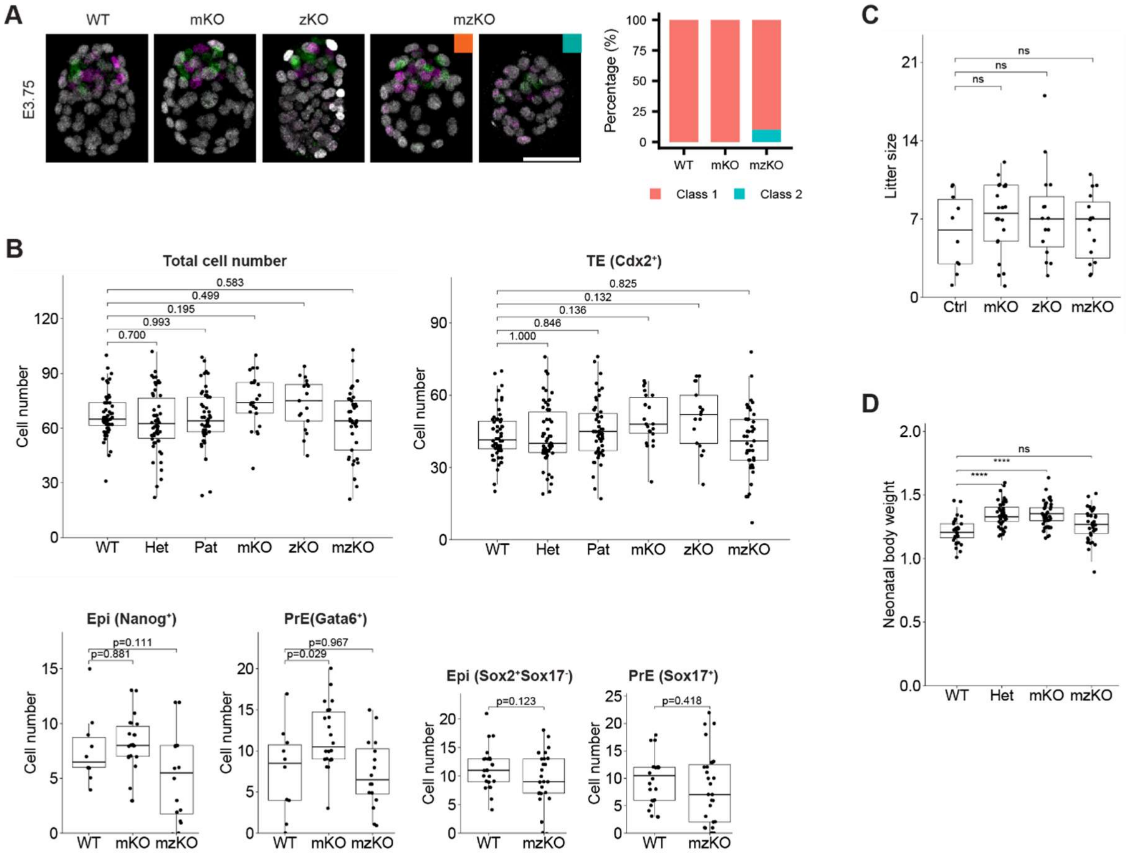


**Figure S9.** In vivo phenotypes of *Mif* knockout.

**A:** Immunofluorescence of trophectoderm (TE; Cdx2$^+$, white), epiblast (Epi; Nanog$^+$, green), and primitive endoderm (PrE; Gata6$^+$, magenta) of in vivo E3.75 blastocysts dissected from uteri (maximum z-projection). WT (*Mif*$^{+/+}$ ♀ × WT ♂); Het (*Mif*$^{+/-}$ ♀ × WT ♂); mKO (*Mif*$^{-/-}$ ♀ × WT ♂); pKO (WT ♀ × *Mif*$^{-/-}$ ♂); zKO (*Zp3-Cre;Mif*$^{fl/fl}$ ♀ × *Mif*$^{-/-}$ ♂); mzKO (*Mif*$^{-/-}$ ♀ × *Mif*$^{-/-}$ ♂). Class 1 and 2 are with and without cavity, respectively. Note that mzKO slightly inhibited blastocyst formation. Scale bar, 50 µm.

**B:** Cell number. N = 56, 54, 51, 22, 17, and 41 for total and TE; Dunnett's test or two-sided Welch's t-test.

**C:** Litter size. Ctrl (*Mif*$^{fl/fl}$ ♀ × WT ♂). N = 10, 24, 15, and 15; Dunnett's test. Note that litter size is unchanged (19).

**D:** Neonatal pup body weight (g). N = 29, 42, 41, and 36 pubs; Dunnett's test.

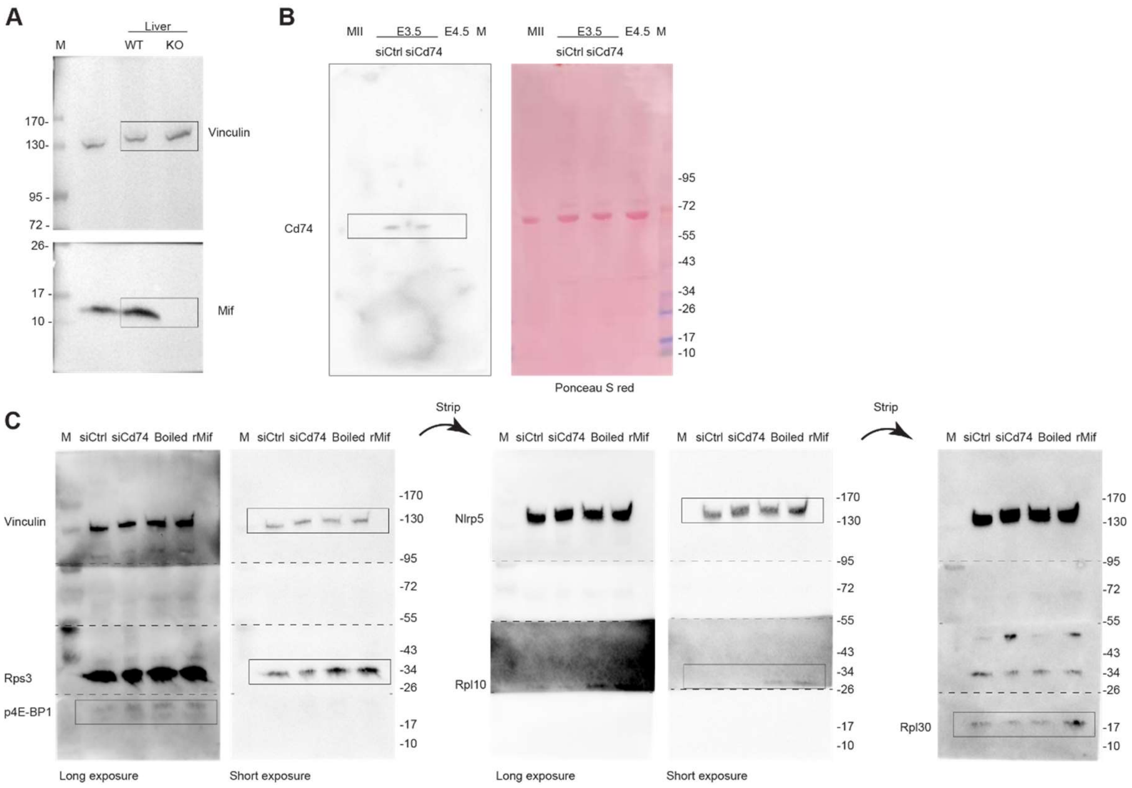


**Figure. S10.** Uncropped Western blot.

**A:** Fig. 1C.

**B:** Fig. 3A.

**C:** Fig. S5D. Spliced membranes (marked by dotted lines) were incubated with antibodies separately and imaged together. M, marker.

**Table S1.** Summary of bioinformatic screen of α-amanitin treated two-cell stage mouse embryos.

| **Class** | **Count** | **Percent** |
|---|---|---|
| 1. Secreted: extracellular annotation | 25 | 0.83 |
| 2. Secreted: signal peptide + extracellular | 28 | 0.93 |
| 3. Secreted: signal peptide | 8 | 0.26 |
| 4. Excluded: secreted + endoplasmic reticulum-localized | 91 | 3.01 |
| 5. Excluded: secreted + transmembrane/lipid-anchor | 53 | 1.75 |
| 6. Candidate: secretion evidence | 4 | 0.13 |
| 7. Candidate: secretion + non-soluble annotation | 1 | 0.03 |
| 8. Not identified as secreted | 2737 | 90.57 |
| UniProt mapped | 2947 | 97.52 |
| Total | 3022 | 100 |

Note: total count of class 1–7 is 210 and of class 1-3 is 61.

**Table S2.** Summary of ELISA results.

| Exp. | Results | | | | | |
|---|---|---|---|---|---|---|
| 1 | *Standard curve recovery (%)* | | | *100.5 ± 1.799* | | |
| | **Condition** | **Stage** | **Density** | **Concentration** | **Mass per embryo** | **N** |
| | MII oocyte | MII | | | 18.8 ± 1.22 | 2 |
| | Medium | | | u.d. | | 2 |
| | WT | E1.5 | 1.25 | 277.3 ± 65.88 | 0.213 ± 0.0584 | 4 |
| | WT | E2.5 | 1.25 | 100.4 ± 18.92 | 0.078 ± 0.0141 | 4 |
| | WT | E3.5 | 1.25 | 365.7 ± 74.11 | 0.324 ± 0.074 | 4 |
| | WT | E4.5 | ~1 | 1417 ± 82.42 | 1.47 ± 0.217 | 4 |
| | zKO | E1.5 | 1.25 | 417 ± 68.33 | 0.302 ± 0.0485 | 4 |
| | zKO | E2.5 | 1.25 | 488.6 ± 288.8 | 0.374 ± 0.215 | 4 |
| | zKO | E3.5 | 1.25 | 426.8 ± 80.72 | 0.339 ± 0.0644 | 4 |
| | zKO | E4.5 | ~1 | 955.4 ± 53.8 | 0.8 ± 0.0589 | 4 |
| 2 | *Standard curve recovery (%)* | | | *99.17 ± 3.102* | | |
| | **Condition** | **Stage** | **Density** | **Concentration** | **Mass per embryo** | **N** |
| | MII oocyte | MII | | | 22 ± 1.2 | 2 |
| | mKO | E1.5 | 1.25 | u.d. | u.d. | 3 |
| | mKO | E2.5 | 1.25 | u.d. | u.d. | 3 |
| | mKO | E3.5 | ~1 | u.d. | u.d. | 3 |
| | mKO | E4.5 | ~1 | u.d. | u.d. | 3 |
| | mzKO | E1.5 | 1.25 | u.d. | u.d. | 2 |
| | mzKO | E2.5 | ~1 | u.d. | u.d. | 2 |
| | mzKO | E3.5 | ~1 | u.d. | u.d. | 2 |
| | mzKO | E4.5 | ~1 | u.d. | u.d. | 2 |
| 3 | *Standard curve recovery (%)* | | | *99.82 ± 3.754* | | |
| | **Condition** | **Stage** | **Density** | **Concentration** | **Mass per embryo** | **N** |
| | LD | E3.5–4.5 | 0.033 | 21.93 ± 5.276 | 0.632 ± 0.154 | 3 |
| | HD | E3.5–4.5 | 1 | 1090 ± 139.3 | 0.679 ± 0.125 | 2 |
| 4 | *Standard curve recovery (%)* | | | *105.4 ± 3.715* | | |
| | **Condition** | **Stage** | **Density** | **Concentration** | **Mass per embryo** | **N** |
| | Medium | | | 35.82, u.d. | | 2 |
| | WT | E1.5 | 2.5 | 1586 ± 77.58 | 0.599 ± 0.0293 | 2 |
| | WT | E2.5 | ~2 | 207.9, u.d. | 0.0906, u.d. | 2 |
| | WT | E3.5 | ~1.5 | 177, u.d. | 0.0885, u.d. | 2 |
| | WT | E4.5 | ~1.25 | 3347 ± 3.747 | 2.32 ± 0.00259 | 2 |
| | WT | E1.5 | 5 | 1458 ± 276.9 | 0.63 ± 0.12 | 2 |
| | WT | E2.5 | ~4 | 422.9 ± 110.9 | 0.211 ± 0.0554 | 2 |
| | WT | E3.5 | ~4 | 447.9 ± 85.8 | 0.252 ± 0.0483 | 2 |
| | WT | E4.5 | ~3 | 4182 ± 479.9 | 3.31 ± 0.38 | 2 |

Note: data in mean ± s.e.m. from four experiments (exp.) in the following units, density ($\mu l^{-1}$), extracellular Mif concentration ($pg\ ml^{-1}$), Mif mass released per embryo (pg); u.d., undetectable.

**Table S3.** Genetic composition of knockout.

| **Embryo condition** | **Mother genotype** | **Father genotype** | **Embryo genotype** | **Maternal protein** | **Zygotic protein** |
|---|---|---|---|---|---|
| WT | $Mif^{+/+}$ | $Mif^{+/+}$ | $Mif^{+/+}$ | Yes | Yes |
| mKO | $Mif^{-/-}$ | $Mif^{+/+}$ | $Mif^{+/-}$ | No | Yes |
| pKO | $Mif^{+/+}$ | $Mif^{-/-}$ | $Mif^{+/-}$ | Yes | Yes |
| zKO | *Zp3-Cre*;$Mif^{fl/fl}$ | $Mif^{-/-}$ | $Mif^{-/-}$ | Yes | No |
| mzKO | $Mif^{-/-}$ | $Mif^{-/-}$ | $Mif^{-/-}$ | No | No |
